# Discovering Plastic-Binding Peptides with Favorable Affinity, Water Solubility, and Binding Specificity Through Deep Learning and Biophysical Modeling

**DOI:** 10.64898/2026.03.30.715295

**Authors:** Tianhong Tan, Michael T. Bergman, Carol K. Hall, Fengqi You

## Abstract

Microplastic (MP) pollution, which is present in the ecosystem in vast quantities, adversely affects human health and the environment, making it imperative to develop methods for its mitigation. The challenge of detecting or capturing MPs could potentially be addressed using plastic-binding peptides (PBPs). The ideal PBP for MP remediation would not only bind strongly to plastic, but also have other properties such as high solubility in water or great binding specificity to a certain plastic. However, the scarcity or absence of known PBPs for common plastics along with the lack of methods that can discover PBPs with all of the desired properties precludes the development of peptide-based MP remediation strategies. In this study, we discovered short linear PBPs with high predicted water solubility and binding specificity by employing an in-silico discovery pipeline that combines deep learning and biophysical modeling. First, a long short-term memory (LSTM) network was trained on biophysical modeling data to predict peptide affinity to plastic. High affinity peptides were generated by pairing the trained LSTM with a Monte Carlo tree search (MCTS) algorithm. Molecular dynamics (MD) simulations showed that the PBPs discovered for polyethylene, the most common plastic, had 15% lower binding free energy than PBPs obtained using biophysical modeling alone. PBPs with both high affinity and high predicted solubility in water were found by including the CamSol solubility score in the MCTS peptide scoring function, increasing the average solubility score from 0.2 to 0.9, while only minimally decreasing affinity for polyethylene. The framework also discovered peptides with high binding specificity between polystyrene and polyethylene, two major constituents of MP pollution, using a competitive MCTS approach that optimized the difference in affinity between the two plastics. MD simulations showed that competitive MCTS increased the binding specificity of PBPs for polystyrene and identified peptides with relatively great preference for either of the two plastics. The framework can readily be applied to design PBPs for other types of plastic. Overall, the high-affinity PBPs with desirable properties discovered by marrying artificial intelligence and biophysics can be valuable for remediating MP pollution and protecting the health of humans and the environment.

## Introduction

There is a pressing need to remediate microplastic (MP) pollution^1^, which is defined to be plastic waste smaller than 5 mm in size. Enormous quantities of MPs are in the environment; tens of millions of tons are predicted to be in the top 200 meters of the Atlantic Ocean alone^2^. MP pollution is also in lakes^3^, rivers^4^, polar ice caps^5^, soil^6^, and the air^7^. MPs are unintentionally consumed by many organisms^8–10^ including humans^11^. It is expected that MP consumption will negatively impact the health of both individuals and ecosystems^12^ as the concentration of MPs increases^13^. Given the continuous influx of MPs into the environment from direct discharge and from the gradual fragmentation of larger plastic debris^14^, it is imperative to develop effective techniques to detect and eradicate MP pollution.

Plastic-binding peptides (PBPs) may be useful for remediating MP pollution. The key insight behind this thinking is that peptides naturally bind strongly to many materials, including plastics^15–17^. PBPs that bind strongly to MPs can be used to detect pollution, filter MPs from water, or accelerate biodegradation by helping plastic-degrading microorganisms or enzymes adhere to MPs. Recent studies have demonstrated the feasibility of using PBPs in these domains^18,19^. However, there are no known PBPs for many plastics. Having a method that discovers PBPs would enable realization of MP remediation strategies that can protect human health and the environment.

The challenge of discovering PBPs to remediate MP pollution requires new computational tools. The peptides must have high affinity to plastic, be soluble in water (bodies of water are a common sink for MP pollution), and ideally bind preferentially to the target plastic more strongly than other plastics so that MP pollution can be separated into its components. Existing deep learning (DL) peptide discovery methods cannot meet all of these demands. Most DL peptide discovery methods focus on healthcare applications, such as the design of antimicrobial peptides, anticancer peptides, or protein-binding peptides^20–26^. They are not suitable for discovering PBPs for two reasons. First, they may use problem-specific metrics to evaluate peptides, like those used to discover peptides that bind to the major histocompatibility complexes^24^. Second, the methods rely heavily on experimental data, such as the ADP3 database^27^ to discover antimicrobial peptides^21,28^, the CPPsite 2.0 database^29^ to design cell-penetrating peptides^30,31^, or the RCSB^32^ or PepBDB^33^ databases to discover peptide drugs^34^. Experimental data is scarce for PBPs, and all existing data comes from screening peptide libraries. While library screening data can be used to train models that give binary predictions of whether a peptide binds to a material like polystyrene^35^ or gold^36^, the data is qualitative and thus poorly suited for quantitatively optimizing peptide affinity. Others have discovered solid-binding peptides by combining small quantitative experimental datasets with molecular dynamics (MD) simulations^37,38^. However, these methods may require modification for different materials, and may not discover high affinity peptides since they only sample a small fraction of the possible peptide sequences. Overall, existing tools cannot meet the unique demands of PBP discovery. A new approach is required.

We hypothesize that an effective method for discovering PBPs is to combine biophysics-based computational modeling with deep learning (DL). PBPs were recently discovered using the biophysics program PepBD^39^, which uses Metropolis Monte Carlo to sample different amino acid sequences and conformations. Peptides are scored^40^ based on the MM/GBSA^41^ binding energy which captures the strength of interactions between the peptide and the plastic, and the stability of the peptide in the adsorbed conformation. While PepBD evaluated millions of peptide sequences while searching for plastic-binding peptides, this is still a vanishingly small fraction of possible sequences. High affinity peptides likely remain undiscovered. Moreover, the Monte Carlo methods of PepBD do not use past results to guide future sampling, meaning PepBD cannot learn from past designs. This limitation can be filled by DL models which can learn patterns in PepBD data and intelligently navigate the enormous number of possible peptide sequences. The flexibility of DL models also makes it straightforward to simultaneously optimize peptide affinity with other desirable peptide properties, such as peptide solubility and binding selectivity. This feat is not straightforward using PepBD or other extant computational tools.

In this work, DL and biophysical modeling were combined to discover PBPs that have high affinity to plastic, high predicted solubility in water, and optimized binding specificity for the target plastic. A Monte Carlo tree search (MCTS) algorithm, tailored to identify peptides with high affinity to plastic, was integrated with a long short-term memory (LSTM) score-predictor trained on PepBD data (Figure S1). MD simulations showed that the PBPs found for polyethylene had affinity equal to the best PBPs found previously by PepBD. PBPs with high solubility in water were discovered by adding a solubility term into the evaluation of peptides by MCTS. PBPs with enhanced binding specificity for polystyrene over polyethylene were found by optimizing the affinity difference between the two plastics. The flexibility of the framework to search for PBPs with high affinity, solubility, and binding specificity cannot be readily achieved with existing, state-of-the-art peptide discovery algorithms. The PBPs identified in this work could be instrumental in cleaning up two major components of MP waste – polyethylene and polystyrene. Our method can readily discover PBPs for other common plastics, which can help develop new peptide-based biotechnology tools for combatting MP pollution.

## Results

### Driving PBP discovery with a DL pipeline integrating PepBD, LSTM, and MCTS

Our DL pipeline effectively discovers peptides with high predicted affinity to polyethylene. The pipeline combines a DL model (LSTM) that predicts peptide affinity for polyethylene with a reinforcement learning algorithm, Monte Carlo Tree Search (MCTS), to explore amino acid sequences with high affinity. The LSTM was trained on previous PepBD data^39^ to predict the affinity of a peptide based on its amino acid sequence, where the affinity is equated to the PepBD score, which is a sum of the MMGBSA binding energy of the peptide to polyethylene and the peptide’s internal energy in the bound state. The trained LSTM was used by MCTS to learn the policy that generates peptides with optimal PepBD scores. This framework, which we term “DL”, was used to identify 100 peptides that are predicted to have high affinity to polyethylene. Each peptide is 12 amino acids long and excludes proline and cysteine in order to match the peptides in the PepBD dataset. The scores of the DL peptides are concentrated at the low end of the PepBD designs, resulting in an average score of –51. This average is much lower than the average of –26 for the entire PepBD dataset (Figure 1A) and slightly lower than the average for the best 100 PepBD peptides (Figure S1). Note that a more negative score corresponds to greater predicted affinity. PBPs discovered by both PepBD and DL have significantly lower scores than randomly generated sequences, which act as a negative control. It thus appears that the DL method effectively discovers PBPs with high affinity for polyethylene. However, the accuracy of this claim could be influenced by the many tryptophan included in each peptide. As PepBD could include at most 3 tryptophan per peptide, a concern was that the comparison between PepBD and DL PBPs may have been unfair. We thus repeated PBP generation with a three-tryptophan constraint and found that the constraint had little effect on either the average or the best scores of the DL PBPs (Figure 1A). Therefore, DL effectively discovers peptides with high predicted affinity to polyethylene.

**Figure 1.**
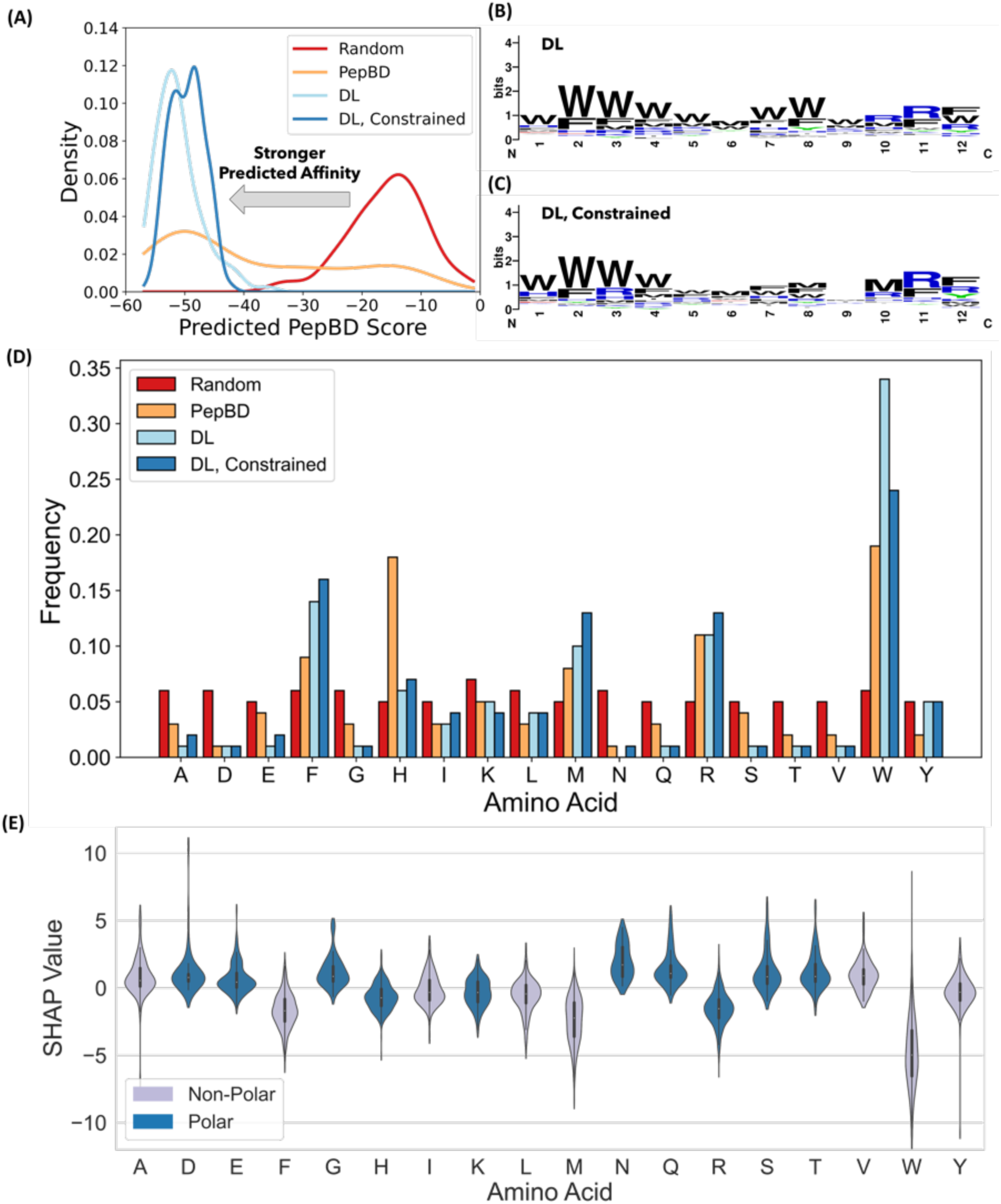
DL discovers PBPs with scores equal to the best PBPs found by PepBD, where the PBPs are enriched in amino acids with bulky side chains. (A) Comparison of predicted scores for PBPs discovered via random generation of amino acid sequences, DL, DL with the three-tryptophan constraint, and PepBD. The score distribution histogram is fitted with a kernel density estimate, a method for smoothing a data distribution, and is shown for clarity. (B) Sequence logo of PBPs discovered by DL. (C) Sequence logo of PBPs discovered by DL with the three-tryptophan constraint. (D) Amino acid frequency for the four discovery methods in (A). Each dataset includes 100 peptides. (E) SHAP value distributions for amino acids sampled from the PBPs discovered by DL with the three-tryptophan constraint. A total of 500 peptides were evaluated, giving a total of 6,000 amino acids. The violin plots show the distribution of SHAP values for all occurrences of the amino acid. Peptides are colored gray if non-polar and blue if polar.

The high predicted affinity of PBPs found by DL could be traced, in part, to enrichment in amino acids that are either hydrophobic or have bulky side chains. While sequence logos^42^ do not show a dominant pattern or motif in the amino acid sequence, the top-scoring PBPs have many tryptophan (W) residues, particularly at the peptide N-terminus (Figure 1B and C). Other residues that appear frequently are hydrophobic, such as phenylalanine (F) and methionine (M), or have bulky side chains, such as arginine (R) and histidine (H) (Figure 1D). In contrast, amino acids with hydrophilic side chains, such as glutamine (Q) and asparagine (N), or small side chains, such as alanine (A) and serine (S), are uncommon in the discovered PBPs. The cationic residues, arginine (R) and lysine (K), occur much more frequently than the anionic residues glutamic acid (E) or aspartic acid (D). We explore why the designs have these features in the following section.

### Interpreting the contributions of amino acids to peptide affinity

To understand the relationship between the peptide score and the amino acid sequence, the SHapley Additive exPlanation (SHAP) values were calculated for all amino acids. For PBP design, a SHAP value^43^ represents how an amino acid contributes towards the PepBD score, meaning that an amino acid decreases the PepBD score and increases the predicted affinity when its SHAP value decreases. The relative importance of each amino acid to the predicted affinity to polyethylene can be elucidated by comparing their SHAP values. Using the PBPs found with DL and the three-tryptophan constraint, we calculated the SHAP values for all amino acids averaged over all residues (Figure 1E) and the position-specific SHAP values (Figure S2). The corresponding information was also calculated for the top 100 PepBD PBPs (Figure S3). Comparison of Figure 1D and Figure 1E shows that the amino acid frequency generally increases as the average SHAP value decreases. Average SHAP values are negative for residues with hydrophobic or bulky side chains, including phenylalanine (F), histidine (H), methionine (M), arginine (R), and tryptophan (W), while average SHAP values are positive for amino acids with small side chains, including alanine (A), glycine (G), or aspartic acid (D). These trends are consistent with the following physical explanation for peptide adsorption to plastic surfaces. Since polyethylene is non-polar, the dominant interactions in peptide adsorption are van der Waals forces and the solvent energy at the plastic surface. A bulky amino acid side chain, like the indole group of tryptophan, can form strong van der Waals interactions with polyethylene and reduce the area of the plastic-water interface. Both of these properties are favorable, so there is a driving force to incorporate amino acids with bulky side chains into PBPs. This is reflected in PBP designs having large masses relative to random amino acid sequences of the same length (Figure S4). This can also explain why lysine (K) and arginine (R) are more common in the designed PBPs than aspartic acid (D) glutamic acid (E), despite each residue having a net charge. In addition to size, hydrophilicity of the side chain matters. Hydrophobic residues aid binding to polyethylene as their unfavorable interactions with water promote adsorption. However, some residues must be exposed to the solvent, and hydrophilic residues are most favorable at these locations. This explains why PBPs still contain a significant fraction of hydrophilic residues. A third factor to consider is that the contribution of an amino acid to peptide adsorption depends on the entire amino acid sequence, not the isolated amino acid. This is clearly demonstrated by the SHAP values spanning positive and negative values for all amino acids, indicating that every amino acid can either improve or worsen affinity depending on what the rest of the amino acid sequence is.

### Designing PBPs with both high affinity and water solubility

We next aimed to discover PBPs that have both affinity to polyethylene and high solubility in water. PBPs require solubility in water if they are to remediate MP in aqueous environments, since low solubility would limit the capacity for capturing or detecting MPs. One way to discover PBPs with both high affinity and solubility in water is to constrain the number of hydrophobic residues. The three-tryptophan constraint previously used is an example of such a strategy. However, this solution is not optimal because reducing the frequency of tryptophan was accompanied by an increase in the frequency of other hydrophobic residues, such as methionine (M) and phenylalanine (F) (Figure 1D). Additionally, the influence of an amino acid on peptide solubility depends not just on the fraction of hydrophobic residues, but also on how the hydrophobic residues are dispersed in the amino acid sequence.

Our method for increasing water solubility was to introduce a peptide solubility term into the MCTS reward function (see Methods for complete description). The solubility term used is the CamSol solubility score^44^, a method for predicting protein solubility based on the amino acid sequence. We also experimented with predicting peptide solubility using the transmembrane tendency scale^45^ (Figure S5 and Figure S6). The three-tryptophan constraint was not used when including the solubility term in the score function.

The modified DL discovery framework found PBPs for polyethylene with both high solubility in water and high predicted affinity, as desired. DL-based peptide discovery was performed with different relative importance assigned to affinity and solubility (Figure 2A). The relative importance was varied by applying a scaling factor (SF) that multiplies the CamSol solubility value (see Methods). When SF is small, affinity dominates solubility and the amino acid composition of PBPs matches designs without considering peptide solubility. When SF is large, solubility dominates affinity and PBPs are enriched in highly soluble amino acids but have poor PepBD scores. Intermediate SF values between 2.0 and 5.0 give rise to PBPs with both high solubility and low PepBD scores. These PBPs have a much lower frequency of tryptophan (W) compared to the original DL PBPs. This was accompanied by an increase in the frequency of charged residues like arginine (R) and lysine (K). This is in stark contrast to the tryptophan constraint, where the lower frequency of W was accompanied by an increased frequency of other hydrophobic residues (see Figure 2C). It is interesting to note that MCTS balances affinity and solubility by making PBPs amphiphilic, where hydrophilic and hydrophobic residues are concentrated at the N– and C-termini, respectively (Figure 2B). Amphiphilicity was not present in the PepBD data and appears to have been discovered by MCTS.

**Figure 2.**
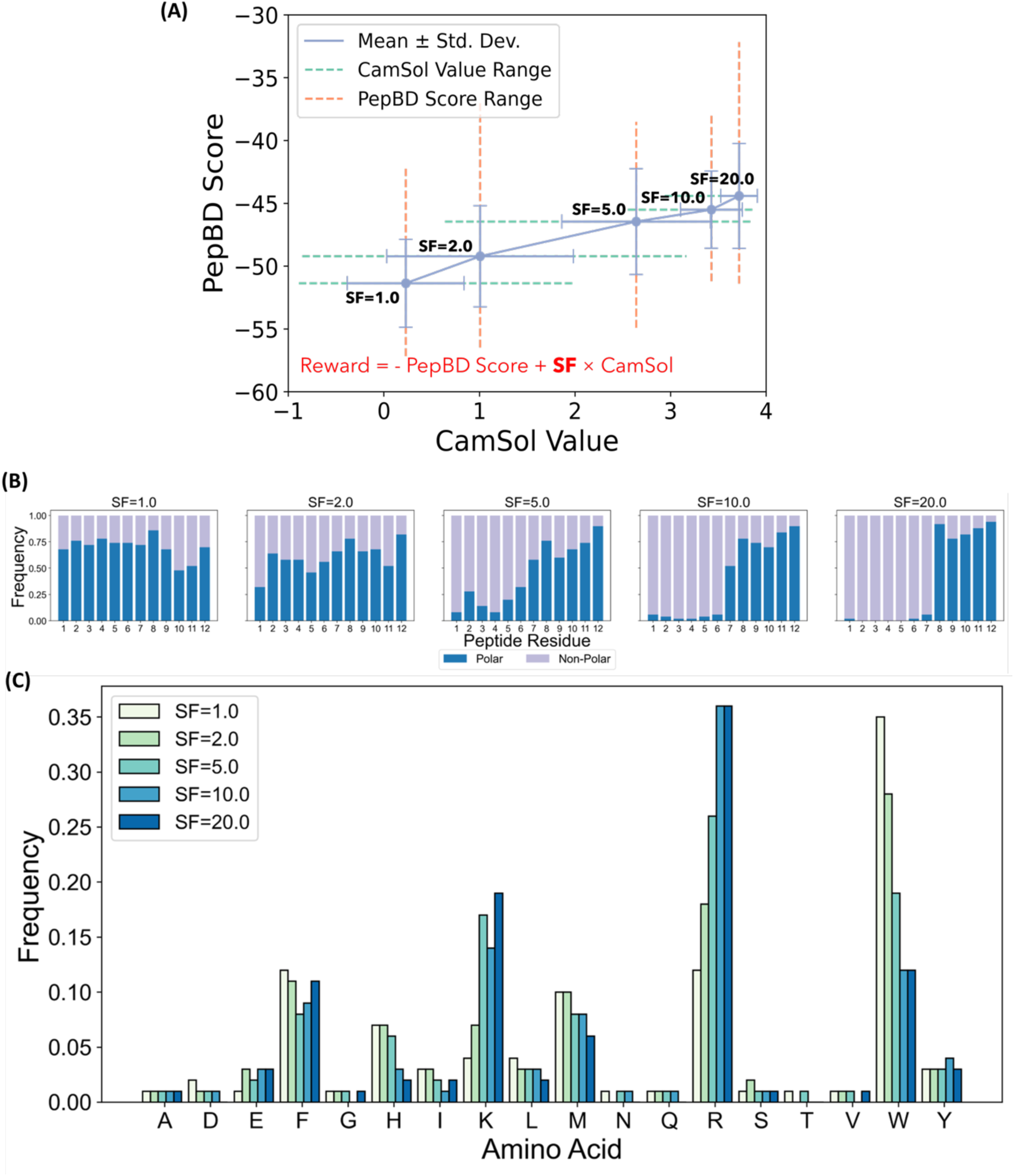
DL-based discovery can simultaneously maximize PBP affinity and water solubility. (A) Dependence of the CamSol and PepBD score of DL-discovered PBP with different scaling factors (SF) that control the relative importance of the PepBD score and CamSol value in the MCTS reward function. Standard deviations and ranges of each value are provided. The MCTS reward function is shown in red text. (B) Fraction of polar and non-polar residues at reach peptide residue for SF values set to 1.0, 2.0, 5.0, 10.0, and 20.0. Polar residues are defined as arginine, histidine, lysine, aspartic acid, glutamic acid, serine, threonine, asparagine, and glutamine. Non-polar are defined as alanine, isoleucine, leucine, methionine, phenylalanine, tryptophan, tyrosine, and valine. (C) Amino acid frequencies at all five SF values. Sequence logos for all SF values are provided in Figure S7.

### Validating the high affinity of DL-discovered PBPs for polyethylene in MD simulations

Calculations of binding free energy (Δ*G*) in molecular dynamics (MD) simulations indicate that the DL PBPs found either with the three-tryptophan constraint or solubility term have as good or greater affinity to polyethylene than PepBD PBPs. Δ*G* was evaluated for 12 DL PBPs with the three-tryptophan constraint, 12 DL PBPs including CamSol (DL + CamSol) with SF set to 2.0, 12 randomly generated amino acid sequences, and 20 best PepBD PBPs from our previous work^39^ (Figure 3). The PBPs discovered by PepBD and the two DL methods have an average Δ*G* much lower than that of the random sequences, indicating that the methods do much better than random chance at finding PBPs as hoped. The DL + CamSol PBPs have a slightly weaker average Δ*G* than DL PBPs (–23.8 vs. –27.0 kcal/mol), mirroring the small decrease in the average PepBD score when adding the CamSol term to the MCTS reward function (Figure 2A). The slight decrease in affinity upon introducing CamSol into the score function is balanced by a significant increase in the CamSol value (i.e., the predicted solubility). The average Δ*G* of DL + CamSol PBPs is equivalent to the average Δ*G* of PepBD PBPs (–23.5 kcal/mol). Thus, MD simulations indicate that DL discovers PBPs with high affinity to polyethylene, while DL + CamSol discovers peptides with both high predicted water solubility and affinity to polyethylene.

**Figure 3.**
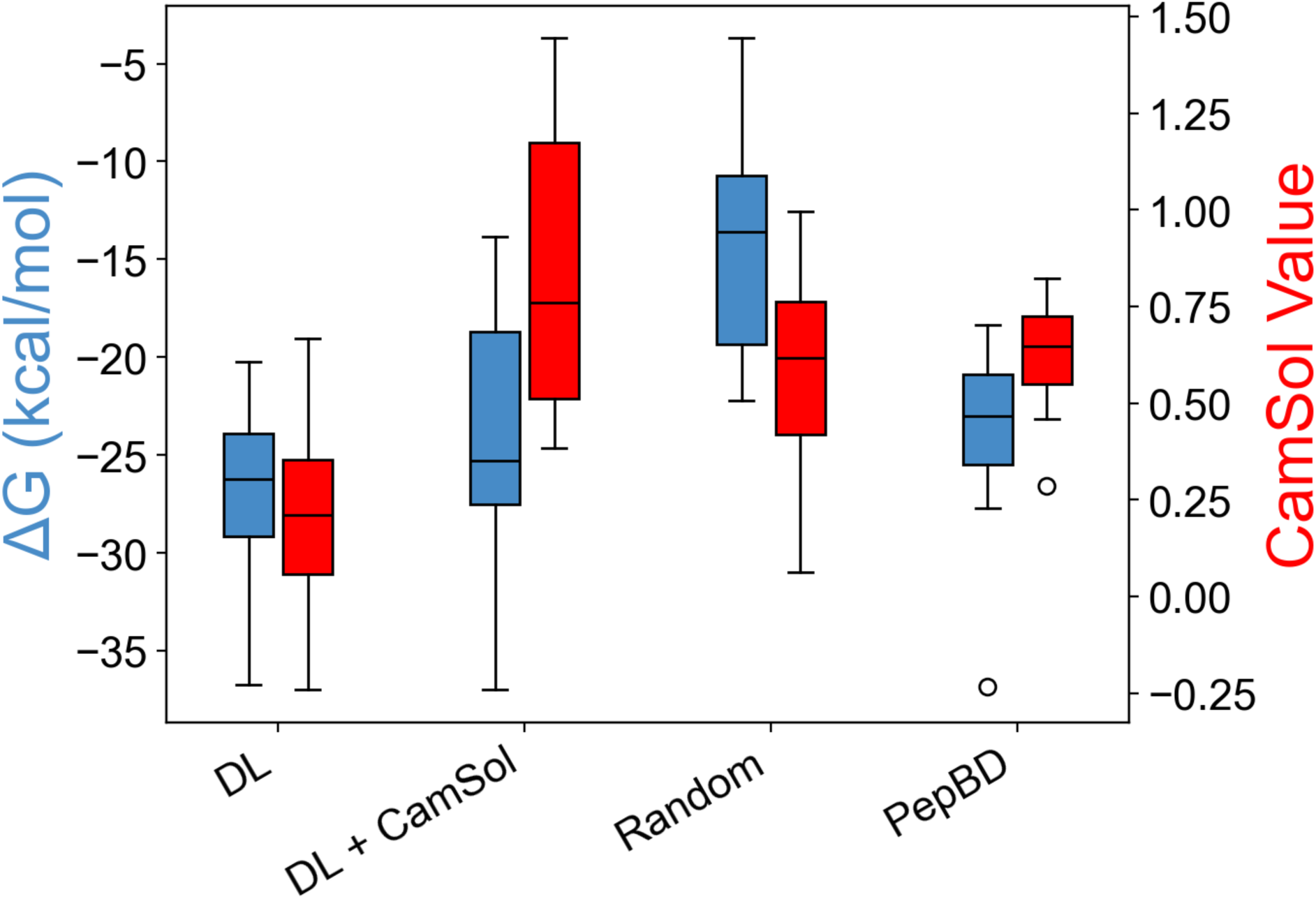
MD simulations show that DL discovers PBPs with equal or higher affinity for polyethylene than PBPs from PepBD, and that peptide solubility can be increased without decreasing affinity. Results are shown for peptides discovered using DL with the three-tryptophan constraint (DL), DL including CamSol with SF set to 2.0 (DL + CamSol), randomly generated amino acid sequences (Random), and PepBD. 20 PepBD PBPs were evaluated, and 12 peptides were evaluated for the other discovery types. Binding free energies (Δ*G*) are shown in blue, and CamSol scores are shown in red.

### Optimizing specificity of PBPs between polyethylene and polystyrene

We now shift our focus to the discovery of PBPs that bind specifically to either polyethylene or polystyrene. This is of interest because polystyrene is a common component of MP waste, and PBPs that discriminate between common plastics could help identify or separate these different components of MP pollution. Can short linear PBPs discriminate between similar plastics like polyethylene and polystyrene? It might be possible given the ability of proteins to discriminate between different types of polymers^46^ and the different composition of proteins adsorbed to plastic nanoparticles with different surface properties^16^. We thus investigate whether we can discover PBPs that show significant preference for polystyrene over polyethylene, and vice versa. This is a challenging task as both polyethylene and polystyrene are simple, aliphatic polymers.

The DL framework was easily modified to search for peptides with large affinity differences between polyethylene and polystyrene. Two separate LSTM score predictors were trained: one that predicts PepBD scores for polyethylene, and another that predicts PepBD scores for polystyrene. Both LSTM models were trained on PepBD data previously generated^39^. The MCTS reward function was modified to maximize both the CamSol score and the difference in the PepBD scores between polyethylene and polystyrene. This latter feature means peptides have favorable scores if they are predicted to have a large affinity difference between the two plastics. We term this approach “competitive” discovery to distinguish it from the previous method used in this paper which we will term “non-competitive” discovery.

Competitive discovery identified peptides with large predicted PepBD score differences between polyethylene and polystyrene; this was reflected to a lesser degree in MD simulations. We searched for fifty peptides that bind specifically to polyethylene over polystyrene, and fifty peptides that bind specifically to polystyrene over polyethylene. The score distributions show the desired large gap in the PepBD scores for the two plastics using both discovery methods (Figure 4A). A side effect of optimizing the score difference was a moderate decrease in the PepBD scores compared to non-competitive discovery. Evaluating the binding enthalpies (Δ*H*) and free energies (Δ*G*) in MD simulations of these PBPs to both polystyrene and polyethylene, however, shows that nearly all PBPs bind more strongly to polyethylene than polystyrene (Figure 4B, Figure S8). The greater affinity for polyethylene over polystyrene is correlated with stronger Lennard Jones interactions between the peptide and polyethylene (Figure S9). We believe this is attributable to the amorphous polystyrene surface having greater roughness than the crystalline polyethylene surface. While all peptides are predicted to preferentially bind polyethylene, competitive design still appears to have improved PBP specificity. The average Δ*H* difference between polystyrene and polyethylene is 8 kcal/mol lower for competitive design compared to non-competitive polystyrene PBPs. Competition found peptides with a larger difference in Δ*G* between polyethylene and polystyrene relative to non-competitive designs, especially for polystyrene (Table 1). Thus, competitive design was moderately successful in finding specific PBPs with enhanced binding specificity to the target plastic.

**Figure 4.**
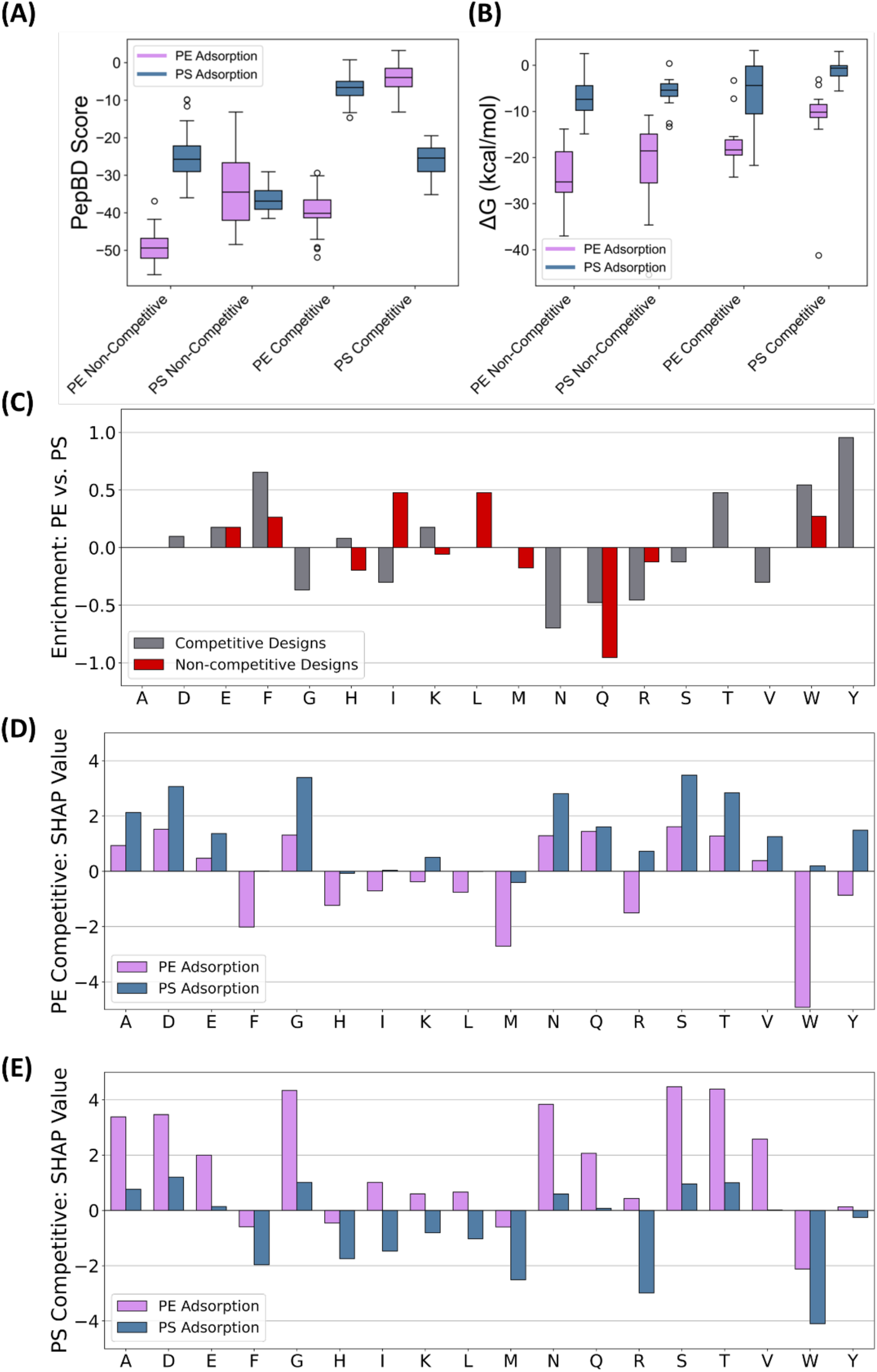
Adding “competition” increases preference of discovered PBP to polystyrene (PS) over polyethylene (PE) and alters amino acid composition. (A) Predicted PepBD scores for competitive and non-competitive DL discovery of peptides that bind to polyethylene (purple) and polystyrene (blue). (B) Binding free energies for competitive and non-competitive DL discovery of peptides that bind to polyethylene (purple) and polystyrene (blue) evaluated in MD simulations. (C) Enrichment of amino acids in polyethylene-specific peptides relative to polystyrene-specific peptides for both competitive (grey) and non-competitive discovery (red). (D) SHAP value for each amino acid type competitive discovery of peptides that bind specifically to polyethylene. Scores are shown for the sequences in binding to either polyethylene (purple) or polystyrene (blue). (E) SHAP value for each amino acid type competitive discovery of peptides that bind specifically to polystyrene. Scores are shown for the sequences in binding to either polyethylene (purple) or polystyrene (blue).

**Table 1.**
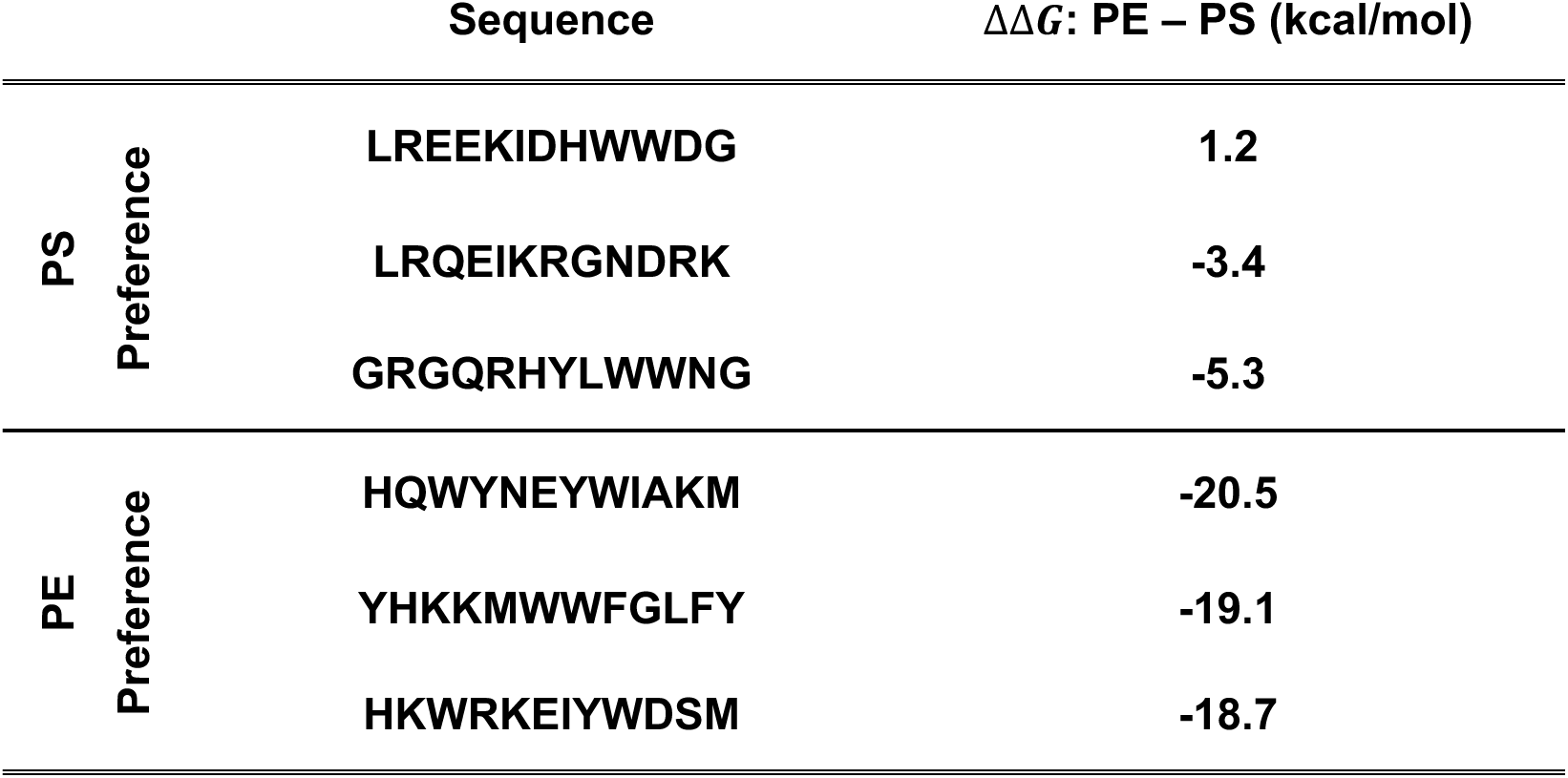
Peptides with substantial differences in binding energies to polystyrene or polyethylene. Three peptides are provided that have greater than average preference for polyethylene (PE) or for polystyrene (PS). For each peptide, the sequence and binding free energy difference, Δ*G_PE_* − Δ*G_PS_*, are listed. For reference, the average ΔΔ*G* without competitive design was –16.5 kcal/mol for PE and –15.5 kcal/mol for PS.

| | Sequence | $\Delta\Delta G$: PE – PS (kcal/mol) |
| --- | --- | --- |
| PS<br>Preference | LREEKIDHWWDG | 1.2 |
|  | LRQEIKRGNDRK | -3.4 |
|  | GRGQRHYLWWNG | -5.3 |
| PE<br>Preference | HQWYNEYWIAKM | -20.5 |
|  | YHKKMWWFGLFY | -19.1 |
|  | HKWRKEIYWDSM | -18.7 |

Competitive discovery improved binding specificity to the target plastic by enriching the peptide in amino acids that prefer one plastic over the other. To reach this conclusion, we first calculated the amino acid enrichment from the amino acid frequencies (Figure S10). Enrichment is defined as

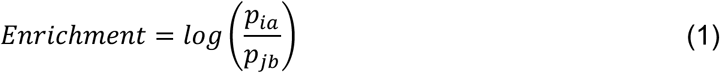

where *p_ia_* is the fraction of amino acid type *i* in peptides discovered using method *a*. The amino acid enrichment differs significantly between competitive and non-competitive discovery (Figure 4C). Polyethylene-specific peptides were enriched in phenylalanine (F), tryptophan (W), and tyrosine (Y), while polystyrene-specific peptides were enriched in arginine (R), glutamine (Q), asparagine (N), and isoleucine (I). To explain the enrichment differences between amino acids, a SHAP analysis was performed on polyethylene-specific peptides (Figure 4D) and polystyrene-specific peptides (Figure 4E). We find that there is a correlation between amino acids enrichment and the difference between the SHAP values for the two plastics (Figure S11). For example, W has a larger SHAP value difference between polyethylene and polystyrene for polyethylene-specific peptides (Figure 4D) than for polystyrene-specific peptides (Figure 4E), in other words, W is observed to be enriched in PBPs that are predicted to have specificity to polyethylene. Phrased another way, binding specificity of a PBP to a target plastic increases by using amino acids that are predicted to increase binding affinity to the target plastic much more than the off-target plastic. As the SHAP value depends on the entire sequence, it is important to note that the effect of an amino acid on binding specificity cannot be determined by the isolated amino acid, but rather depends on the full amino acid sequence of the peptide.

## Discussion

We propose a methodology developed to discover short linear PBPs combines biophysical modeling and DL. We train an LSTM trained on biophysical data from PepBD to predict peptide affinity to the common plastics polyethylene and polystyrene. MCTS utilizes the trained LSTM to efficiently generate novel peptide sequences predicted to have high affinity to either plastic. MD simulations show that the best discovered PBPs have slightly greater affinity to polyethylene compared to the best PBPs found using only PepBD. The framework also discovers PBPs with high affinity and other desired properties (e.g. solubility in water and binding specificity) through a highly customizable MCTS reward function.

It is notable that competitive discovery found PBPs with a significantly different binding preferences to polystyrene versus polyethylene. This is contrary to our expectation that peptides would not easily discriminate between the two plastics: they are both hydrophobic and can form neither hydrogen bonds nor ionic interactions, meaning peptide-plastic interactions should be dominated by non-specific van der Waals and hydrophobic interactions. Yet, MD simulations show the difference in Δ*G* and Δ*H* of peptide binding to the two plastics covers a large range (Figure S12). Peptides on opposing ends of this distribution are expected to display different binding preference between polyethylene and polystyrene. The ability of proteins to bind specifically to polymers has been documented by Kumar et al.^46^ and Lundqvist et al.^16^ They found protein adsorption to polystyrene nanoparticles varied with curvature and surface charge of the nanoparticles, but the authors are not aware of such findings for short linear peptides. Experimentally measurements of the affinity difference between polystyrene and polyethylene for these PBPs will be essential to validate our computational predictions.

Combining DL and biophysics is useful for discovering peptides that bind to plastics or other solid materials. The scarcity of quantitative experimental data on peptide-surface interactions makes computational modeling essential for quantitatively exploring why peptides bind strongly to plastics or other materials. DL models can be trained on biophysical modeling data to accelerate exploration of the enormous number of peptide sequences, accelerating PBP discovery. Biophysical evaluation of the peptide designs is essential to verify the efficacy of DL designs. The trained DL models can also find patterns in the modeling data, such as the SHAP analysis in this work, which can guide future peptide discovery and explain why certain peptides bind stronger than others to a given material. The DL framework also permits peptide properties other than affinity to plastic to be optimized during peptide discovery by incorporating ML models trained to predict other peptide properties. While we focused on optimizing peptide solubility and binding specificity, other properties that could be optimized using existing models include peptide self-aggregation^47^, immunogenicity^48^, or toxicity^49^. The overall discovery process (generate PepBD data, train the DL models, and search for peptides) takes only days to complete and does not rely on experimental data. The method developed in this work to discover PBPs can readily be used to discover peptide binders for other materials, such as silica or metals. A major potential bottleneck is an accurate biophysical description of interactions between the peptide and the receptor. Fortunately, models exist for many materials of interest to in biotechnology^50–52^. Different receptors may also require more detailed descriptions of the peptide-receptor complex to accurately predict peptide affinity to the receptor. More potent PBPs might be found with the discovery of advanced large language models (LLMs) such as GPT-4^53^, Gemini^54^, PaLM 2^55^, and LLaMA 2^56^. Alternatively, structural information on the peptide-receptor complex could be captured by using graph neural networks and other novel architectures^57^.

We believe that the PBPs discovered in this work could be broadly helpful in remediating MP pollution. The PBPs can be implemented into sensors to detect MP pollution levels in water, incorporated into water purification filters to remove MP, and engineered into the genomes of plastic-degrading microorganisms^58^ to help the microbes adhere to the plastic to enhance MP decomposition. Peptide-based strategies may be particularly effective for nanometer-sized plastics: PBPs interact with plastics via adsorption, which is driven by surface area, and nanoscale materials often have large surface areas per mass. Having tools suitable for nanoplastics is important since such particles are generally harder to detect and capture than micro– or millimeter sized plastics. The suitability of peptide-based strategies to remediate nanoplastics is suggested by the recent development of a peptide-based biosensor that can detect nanometer-sized polystyrene^19^. We hope that by discovering PBPs with high affinity for two common plastics, enhanced solubility in water, and improved binding selectivity between common components of MP pollution, we can begin to realize the application of peptides towards MP pollution.

## Methods

### Preparation of PepBD dataset

Two datasets of polyethylene-binding and polystyrene-binding peptides were generated using PepBD, as described in previous work^39^. Given a starting peptide sequence and structure, PepBD uses a simulated annealing protocol to sample peptide sequences and conformations to search for peptides with low score. The score for the peptide is calculated using

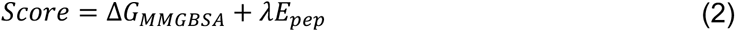

Δ*G_MMGBSA_* is the binding free energy calculated using the MM/GBSA method, *E_pep_* is the internal energy of the peptide, which is included to ensure that the peptide can adopt the bound conformation, and λ is a scaling factor. Both the scores and the peptide sequence are output by PepBD periodically during discovery. The PepBD data for 100 design runs for polyethylene and 55 design runs for polystyrene were collected to generate the dataset used to train the LSTM model as described below. Multiple runs were performed to sample different initial adsorbed conformations. As a sequence could be sampled multiple times and be assigned different scores depending on the peptide structure, only the best (i.e., most negative) score for a sequence was retained. A total of 901,063 sequence: score data points were included in the polyethylene dataset and 405,827 sequence: score data points were included in the polystyrene dataset. PepBD PBPs for polyethylene have a roughly Gaussian distribution of scores with an average of –26 and standard deviation of 11 while PepBD PBPs for polystyrene have a roughly Gaussian distribution of scores with an average of –15 and standard deviation of 9.

### Predicting PepBD score with LSTM

An LSTM model was developed to predict the PepBD score for a peptide sequence (Figure S13). The model consists of an embedding layer followed by three stacked LSTM layers. The embedding layer is used to map integer-encoded amino acids to dense vectors of fixed size. It refines the vector representations so that similar amino acids (in the context of the task) have closer vectors in the embedding space. The embedded representation of the peptide is fed into the first LSTM layer. LSTM is a type of Recurrent Neural Network (RNN) designed to discover patterns in sequences, making it highly suitable for sequential data such as amino acid sequences in peptides^59^. The LSTM layers produce a new sequence of vectors that include higher-level features of the input sequences resulting from interactions between different peptide residues. The last LSTM layer output is fed into a regressor for predicting the PepBD score of the peptide. The training, validation, and test split is 80%, 10%, and 10%, respectively. The Adam optimizer^60^ was used to train the model with a learning rate of 0.001 over a course of 200 epochs. The mean squared error (MSE) between predictions and labels was used as the loss function. The model with the lowest loss on validation set during the training process was selected to evaluate the regression performance on the independent test set (Figure S13).

### MCTS for peptide sequence optimization

A MCTS algorithm^61,62^ was applied to generate peptides with low PepBD scores (Figure 1B). Traditionally harnessed in artificial intelligence for game-play decision-making, MCTS is applied here to navigate peptide sequence space. The reward function is the negative of the PepBD score predicted by the surrogate model. The surrogate model in this work is the trained LSTM. Exploration and exploitation of peptide sequences is balanced using the Upper Confidence Bound applied to the Trees (*UCB1*) formula^63^:

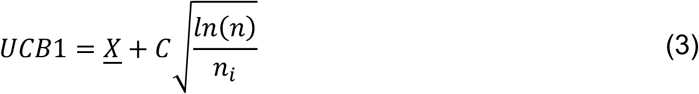

where <u>*X*</u> is the average reward of the node, or the mean outcome of all simulations that have passed through this node; *C* is an exploration parameter which determines the trade-off between exploitation (selecting nodes with high average reward) and exploration (selecting less-visited nodes); *n* is the total number of times the parent node has been visited; and *n_i_* is the number of times the node itself has been visited. The first term promotes nodes with higher average rewards and thus emphasizes exploitation, while the second term promotes nodes that are less frequently visited, especially in the early stages of the search, and thus emphasizes exploration. By combining these two terms, the *UCB1* balances between exploitation and exploration of peptide sequence space. The exploration constant, *C*, tunes this balance: a large value of *C* emphasizes exploration, while a small value emphasizes exploitation.

We now describe the peptide discovery process using MCTS. The process starts with an empty node, then MCTS selects the most promising child node by evaluating *UCB1*, Equation (3), of each possible amino acid. The average reward is calculated using the LSTM surrogate model. Once an action is selected, the algorithm expands the tree by appending an additional amino acid to the decision tree. Next, a simulation or random roll-out is performed, where the new peptide sequence is assessed by using random extensions to gauge its performance. The insights derived from these rollouts guide peptide optimization in subsequent steps. The acquired property values from the simulation are back propagated through the search tree, ensuring that the representation of potential sequences remains up to date. By repeatedly iterating through these four steps, the tree policies are updated to minimize the PepBD score. The optimal peptide sequences, i.e. those with the lowest scores predicted by the surrogate model, are those with the lowest scores encountered during random sequence rollouts (Figure S14). Constrained paths were also explored by pruning certain actions in the action space to refine the search, which was done by forbidding certain actions (appending certain amino acids to the decision tree) when some criterion (i.e., the number of certain amino acids reached a predefined limit) was met. In this study, the constraints were used to ensure that the peptides had desirable properties, e.g. limiting the number of tryptophan to ensure the peptide was soluble in water.

### Simultaneous optimization of peptide affinity and water solubility

In the context of PBP discovery, specific constraints often guide the selection process. Notably, for MP remediation in aqueous environments, the objective includes identifying peptides with a high affinity for binding to a plastic surface, while simultaneously ensuring their solubility in water. We use a multi-objective optimization approach that is compatible with MCTS to tackle this challenge. Solubility is optimized simultaneously with the PepBD score by adding a solubility term to the original MCTS reward function. Two methods for calculating the solubility were tested. The first method estimated peptide solubility using the CamSol method:

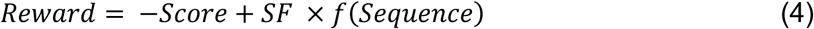

where the peptide solubility is a function, *f*, of the peptide sequence. The second method estimated peptide solubility by summing the transmembrane tendency hydrophobicity (*ttH*)^45^ value of each amino acid:

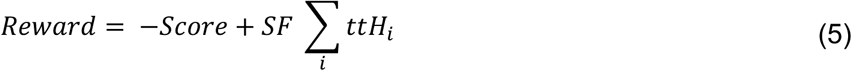

In both equations, *SF* is a scaling factor that controls the relative importance of the solubilty term in the overall peptide score. A variety of *SF* values were evaluated to determine a suitable value that generates peptides with both high affinity and water solubility.

### Optimization of peptide binding preference to one plastic surface over another

To generate the peptides that bind to one plastic surface over another, two separate LSTM models were trained: one predicting the PepBD score for polyethylene, and another predicting the PepBD score for polystyrene. The architecture, hyperparameters, and training process for the polystyrene model are identical with the polyethylene model described in the previous section. Both models have good prediction performance on PepBD scores (polyethylene model: R^2^=0.82, RMSE=4.58; polystyrene model: R^2^=0.81, RMSE=3.90, Figure S13). The reward function was modified by replacing the negative PepBD score for one plastic in the previous approach with the difference between the PepBD scores for polystyrene and polyethylene. The higher this value is, the larger the gap in predicted binding affinity of the peptide for polyethylene versus polystyrene. The CamSol value term was included in the reward function to ensure that the discovered peptides are soluble in water and also do not contain too many tryptophan residues in the sequence. The scaling factor of the CamSol value term was set to be 2.0 based on our earlier experiments because this value provides a good balance between the target to be optimized and the solubility of the peptides. The reward function for generating peptides that have preference for polyethylene over polystyrene is:

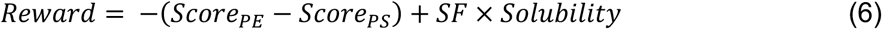

The reward function for generating peptides that have preference for polystyrene over polyethylene is:

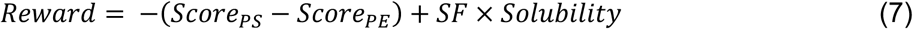

As in equations (4) and (5), *SF* is a scaling factor that controls the relative importance of the solubilty term in the overall peptide score.

### Evaluating peptide affinity using MD simulations

MD simulations were conducted to evaluate the affinity of the discovered peptides for plastic. While enhanced sampling methods like metadynamics^64^ or umbrella sampling^65^ can calculate binding free energies with high-precision, their high computational cost makes them unsuitable for high throughput peptide evaluation as required in this work. Instead, peptides are evaluated in an ensemble of short equilibrium MD simulations to search for the lowest energy bound state. An ensemble of simulations is required because the peptide can bind to the surface in a large number of conformations, and an equilibrium MD simulation at room temperature typically is kinetically trapped in one conformation for long periods of time. A collection of putative, stable adsorbed conformations for each peptide are generated by running a 10 ns simulation at 550K with the peptide placed directly above the surface. The high temperature allows the peptide conformation to change rapidly. To prevent the peptide from diffusing away from the surface due to the high temperature, a harmonic potential using the Upper Wall utility in PLUMED^66^ was added. The harmonic potential pushed the peptide back towards the surface if the distance between the peptide center of mass and the top of the surface exceeded 10 Ā. Once the high temperature simulation completed, k-means clustering was performed with CPPTRAJ^67^ to generate 16 peptide structural clusters. Extracting one representative structure from each cluster gave the starting conformations for the ensemble of simulations. All 16 conformations for each peptide were simulated for 1 ns, then the binding free energy was calculated using Amber’s MMGBSA tool. The 8 simulations with the lowest binding free energy were simulated for an additional 4 ns before again calculating the MMGBSA binding free energy. The lowest binding free energy from the 8 extended simulations represents the most stable conformation and was selected as representative of the peptide’s binding affinity.

Technical details of the simulations are the following. Simulations used the TIP3P water model^68^, GAFF^69^ parameters for plastics with partial charges calculated in our previous work^22^, and the ff14SB force field^70^ for peptides. An extended conformation of the peptide was generated using the tLEaP^71^ module in Amber. Atomistic models of plastic surfaces were taken from our previous work^39^. The peptide was placed on the plastic surface by rotating and translating the peptide such that its long axis was parallel to the plastic surface and the distance between the top of the surface and the peptide center of mass was 4 Å. The system was solvated with tLEaP by adding TIP3P water 15 Å above the peptide and 10 Å below the bottom of the plastic surface. The dimensions of the simulation box parallel to the plastic surface were set to equal the dimensions of the plastic surface. The Amber coordinate and parameter files were converted to Gromacs format using the Parmed^71^ utility of Amber. All simulations were run with Gromacs version 2019.6^72^. For high temperature simulations at 550K, the system was energy minimized using steepest descent for 1,000 steps, heated to 300K in an NVT ensemble over 100 ps, equilibrated at 1 bar and 300K in the NPT ensemble for 200 ps, heated to 550K in the NVT ensemble for 200 ps, then finally running the production phase in the NVT ensemble for an additional 10 ns to generate different peptide bound conformations. After extracting the different bound conformations with k-means clustering, a 100 ps NVT simulation at 300K cooled the system to room temperature, then a production simulation in the NVT ensemble at 300K was run for 1 ns per simulation. The 8 systems with the lowest MMGBSA binding free energy were simulated an additional 4 ns in the NVT ensemble at 300K. Position restraints were applied to all carbons in the plastic using a force constant of 5,000 kJ/mol/nm^2^. The LINCS algorithm^73^ was used to restrain hydrogen positions. Long-range electrostatic interactions were treated using particle mesh Ewald. The time step size was set to 2 fs. NVT and NPT simulations controlled the temperature using the velocity rescaling algorithm^74^ with a time constant of 0.1 ps and separate thermostats for water molecules and the rest of the system. NPT simulations used the semi-isotropic Berendsen barostat^75^ with the x-y dimension changed independently from the z-dimension, a time constant of 5 ps, and an isothermal compressibility of 4.5 × 10^-4^ for all directions.

### Residue level interpretation from SHAP

SHapley Additive exPlanation, or SHAP^76^ was used to understand the contribution of amino acid type and position to PepBD score. SHAP draws on cooperative game theory to evaluate the expected marginal contribution of a feature across all conceivable combinations. SHAP assigns each feature a unique attribution value, indicating its influence on the model’s prediction. For our peptide design problem, a more negative SHAP value attributed to an amino acid indicates that that amino acid contributes more to the plastic binding. The SHAP values of each amino acid type were calculated for peptides extracted using stratified sampling (based on PepBD scores, the number of bins was set to be 10) from both past PepBD PBPs, and PBPs discovered by MCTS. Since the test set included a huge number of peptides, we only extracted a small portion (6,000 amino acids or 500 peptides – the number of peptides discovered with MCTS) using stratified sampling based on the PepBD score of the peptide containing that amino acid for SHAP analysis.

## Data availability

The data presented in this study are available in the manuscript file, the supplementary data file, and source data files in the below GitHub repositories.

## Code availability

Code to run the deep learning model is available in GitHub at https://github.com/PEESEgroup/DL-PBP-Design

The PepBD code used in previous work is available in GitHub at https://github.com/CarolHall-NCSU-CBE/PepBD_Plastics

## Acknowledgements

M.B. acknowledges the Expanse Supercomputing Center allocation provided by ACCESS as well as the Hazel super computing system at North Carolina State University for computational resources needed to conduct molecular dynamics simulations. M.B. was funded from NIH grant 1T32GM133366. F.Y. acknowledges the partial support from the Eric and Wendy Schmidt AI in Science Postdoctoral Fellowship, a Schmidt Sciences program. The authors gratefully acknowledge support from the National Science Foundation (EFMA-2029327) for this work.

## Author Contributions

All authors contributed to project conceptualization, data analysis, and writing of the manuscript. T.T. developed the LSTM and MCTS model and performed the SHAP analysis. M.B. performed molecular dynamics simulations. All authors have given approval for the final version of the manuscript.

