## Supplementary material for "Discovering Plastic-Binding Peptides with Favorable Affinity, Water Solubility, and Binding Specificity Through Deep Learning and Biophysical Modeling": SI Text file

**This PDF file includes:**

- Supplementary Text
- Supplementary Figures 1 to 14

[#]These authors contributed equally to this work

The supplementary materials provide additional data and analyses that augment the findings discussed in the main text. Comprising 14 supplementary figures and accompanying detailed descriptions, these materials are organized to mirror the sequence in which related topics appear in the manuscript, thereby offering a coherent and extended exploration of the study's results. Figure S1 provides an overview of our computational pipeline integrating biophysical modeling and deep learning for PBP discovery. Figure S2 compares the masses of discovered PBPs as well as random PBPs, showing PBPs consistently have large mass relative to random amino acid sequences of the same length. Figures S3 to S5 present additional analyses regarding the discovery of water soluble PBPs. Figures S6 and S7 provide additional analysis on the contribution of amino acids to the predicted affinity to polyethylene. The location specific SHAP values as well as the SHAP value distribution for the amino acids in PBPs discovered by PepBD are illustrated. Figures S8 to S11 encompass the analyses of PBPs discovered using the "competitive" strategy. The MD simulation results, sequence component comparison, and SHAP value correlations are shown to corroborate the discussion in the "Adding competition improves specificity of PBPs for polystyrene over polyethylene" section. Figure S12 compares the Lennard Jones (LJ) interaction energy between peptides and either polyethylene or polystyrene, showing that peptides have stronger LJ interactions with polyethylene than polystyrene for the surface models used in MD simulations. In Figures S13 and S14, information regarding the architecture, training process, and accuracy of the surrogate model as well as the generating process of an optimized sequence is provided. The accompanying text for these figures elaborates on the details of the models developed in our study and ensures the reproducibility of all the computational results. Additional data (i.e., predicted PepBD score, solubility, binding free energy, etc.) regarding the PBPs discovered in this work are also available in a separate data file.

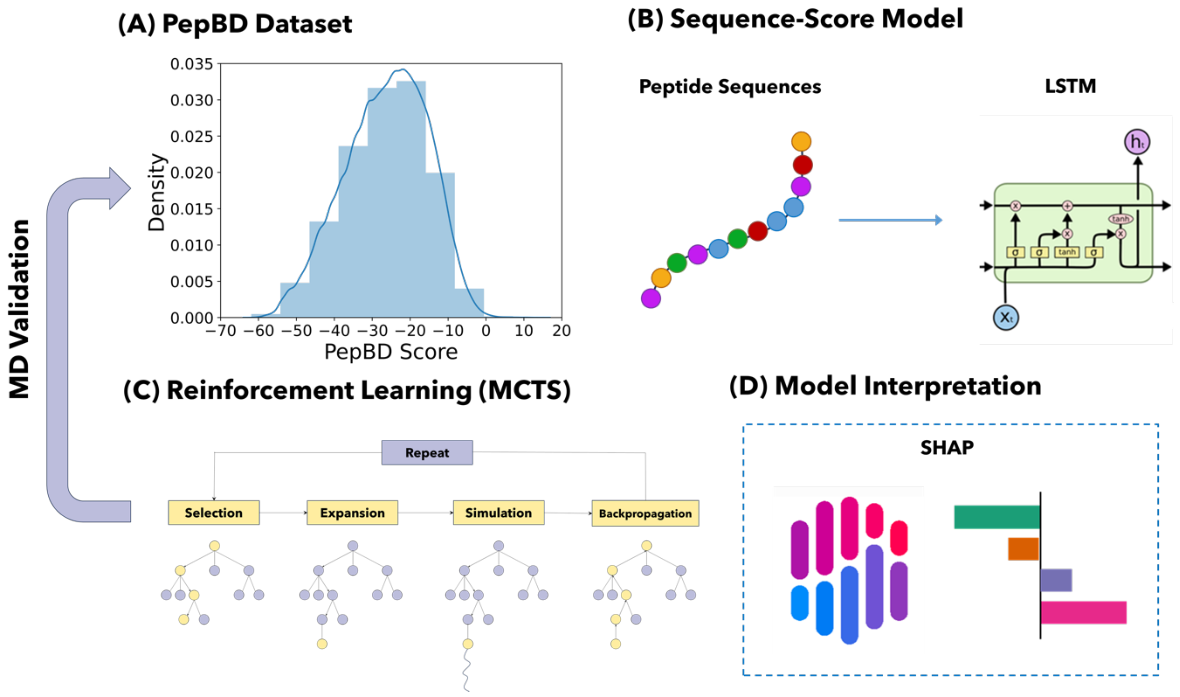


**Figure S1. Overview of modeling framework for discovery of PBPs using DL.** (A) PepBD score distribution of peptides in the polyethylene dataset. (B) LSTM-based model for predicting PepBD score based on peptide sequence information. (C) MCTS algorithm for peptide optimization. One MCTS loop includes steps of selection, expansion, simulation, and backpropagation. (D) SHAP analysis for understanding amino acid contribution to PepBD scores.

Figure S1 presents our novel discovery framework, integrating biophysical modeling with DL, for the development of short linear PBPs. This framework was operationalized by generating two distinct datasets PepBD: one comprising peptides that bind to polyethylene and another for polystyrene binding peptides. The polyethylene dataset encompassed 901,063 sequence-score data points, whereas the polystyrene dataset contained 405,827 data points. The distribution of PepBD scores for polyethylene-binding PBPs approximates a Gaussian curve, with a mean score of -26 and a standard deviation of 11. Similarly, polystyrene-binding PBPs also exhibited a Gaussian score distribution, albeit with a mean of -15 and a standard deviation of 9. Subsequently, an LSTM model was developed to predict the PepBD score based on peptide sequences. This model was further coupled with MCTS to facilitate the generation of optimized peptides characterized by high binding affinity. Furthermore, SHAP analysis was employed to discern the contribution of individual residues within the discovered peptides towards the overall PepBD scores.

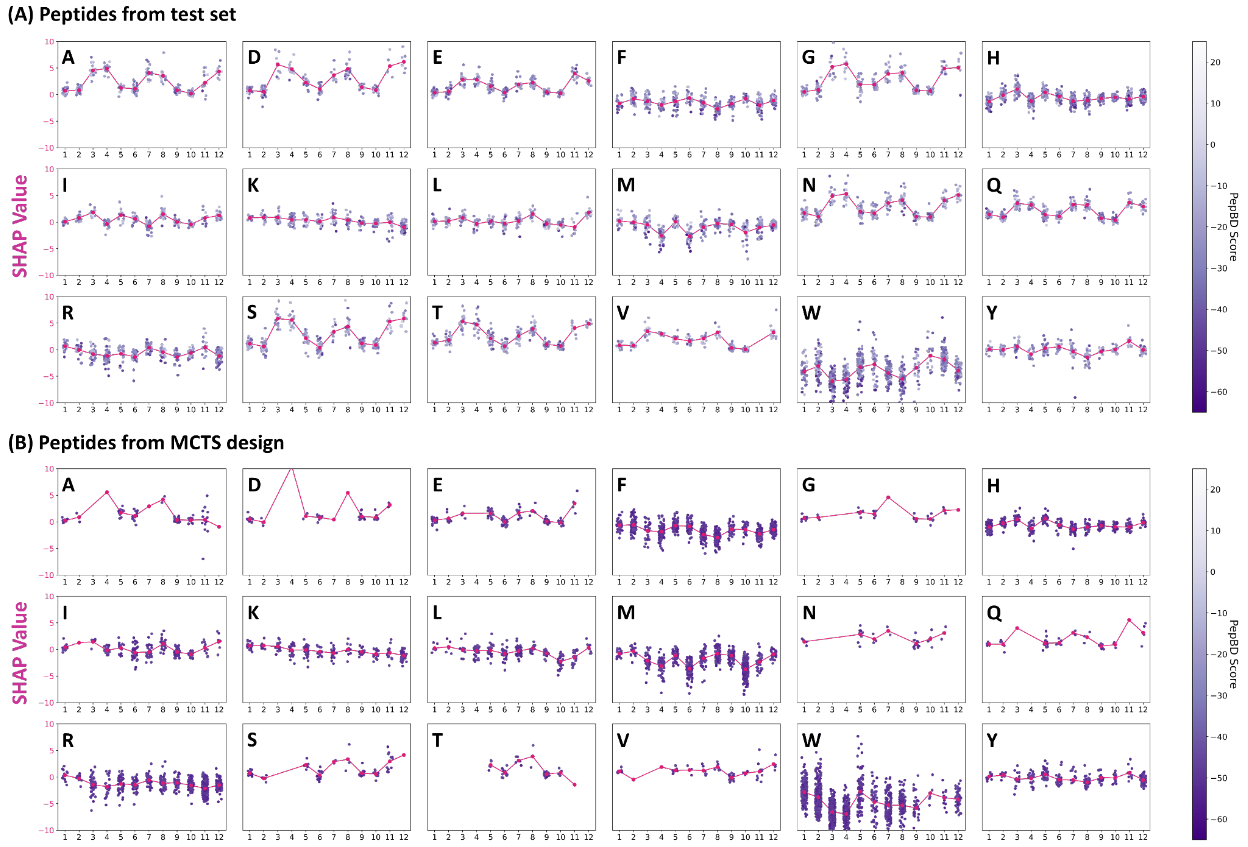


**Figure S2. SHAP values of the amino acids in the peptides at different locations.** (A) Peptides from the PepBD PBPs in the test set. A total of 500 peptides were evaluated using stratified sampling based on PepBD score (the range of PepBD scores was divided into ten categories according to the value, from the minimum to the maximum score, for the purpose of segregation) giving a total of 6,000 amino acids. (B) 500 Peptides from DL discovery.

Figure S2 plots the SHAP values for each amino acid type as a function of position in the peptide. Results are shown both for PBPs from both PepBD and DL. The SHAP value generally does not vary significantly with position, but some residues show some variability, such as tryptophan (W). Two possible explanations for variability of the SHAP value with position is that 1) some amino acids truly are more favorable at certain peptide residues, or 2) this is an artifact arising from the PepBD and DL datasets not exhaustively sampling all possible conformations and sequences. Some residues with highly positive SHAP values at a certain position in PepBD PBPs (e.g., serine (S) at positions 3 and 4 and threonine (T) at position at positions 3 and 4) may have reduced frequency in DL PBPs, which might partially explain why DL PBPs have slightly stronger affinity than PepBD PBPs.

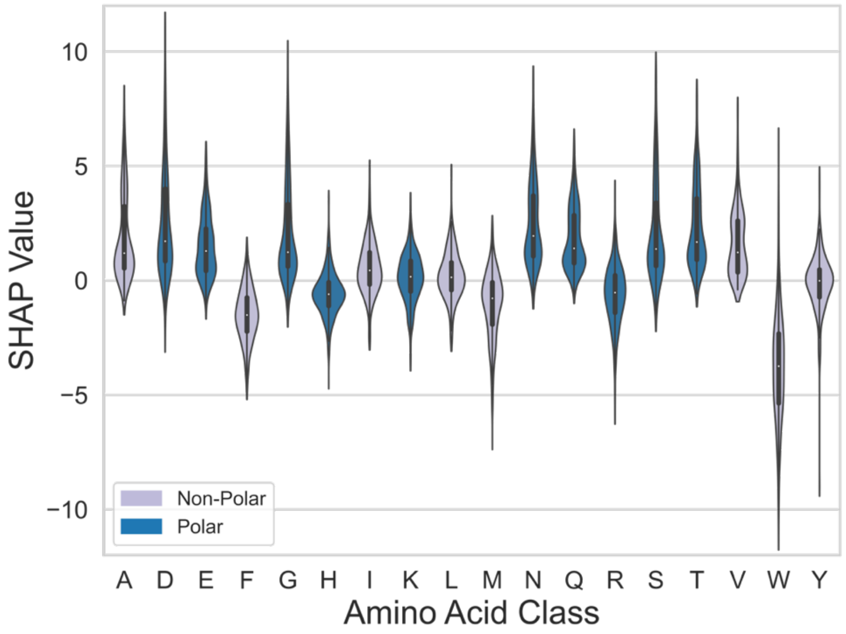


**Figure S3. SHAP value distributions for amino acids sampled from the 500 PepBD PBPs.**

Figure S3 illustrates the distribution of SHAP values derived from an independent test set of 500 data points, originating from PepBD PBPs, analyzed in Figure 1. The analysis reveals that amino acids with positive mean SHAP values in PBPs discovered through DL retain positive mean values in the PepBD dataset, and similarly, amino acids with negative SHAP values maintain their negativity. Moreover, the range of SHAP values observed closely mirrors that of the DL PBPs. This consistency underscores the reliability of SHAP analysis in yielding comparable results for polyethylene-binding PBPs across both DL and PepBD methodologies.

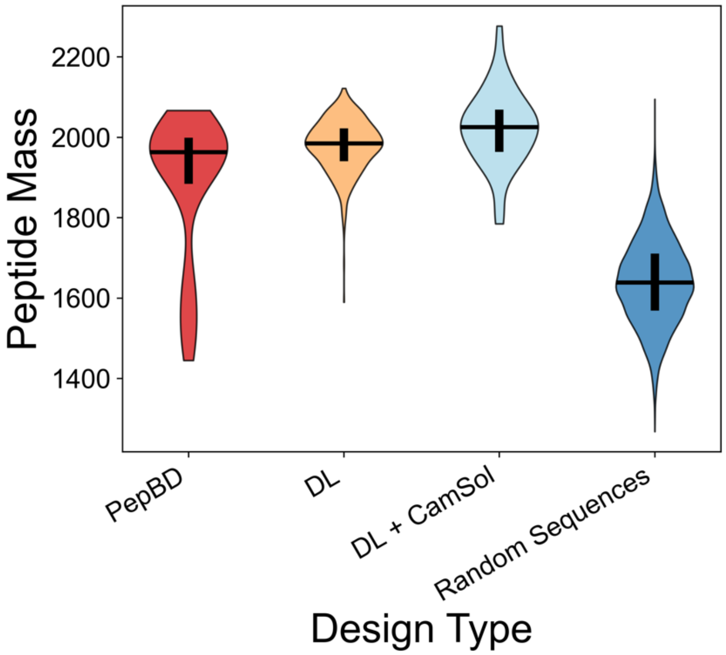


**Figure S4. Peptide masses discovered to bind to polyethylene.** Results are shown for four discovery methods: PepBD, DL with the three tryptophan constraint, DL using CamSol and SF=2.0, and random generation of amino acid sequences. The data for 100 PepBD peptides, 100 DL and DL + CamSol peptides, and 1000 Random sequences are plotted.

Figure S4 shows the distribution of peptide masses for 100 PBPs previously discovered by PepBD, 100 DL PBPs using the constraint of no more than three tryptophan, 100 DL PBPs with the CamSol solubility term and SF=2.0, and 10,000 randomly generated peptide sequences. Random sequences were generated by choosing each of the 20 standard amino acids with equal probability. The PepBD and DL PBPs are all for polyethylene. The average mass for PepBD and DL PBPs are much greater than random peptides, showing the discovery methods enrich for relatively heavy amino acids. Such heavy amino acids can have stronger van der Waals interactions with the plastic and de-solvate a larger area of the surface, both of which should favor peptide binding.

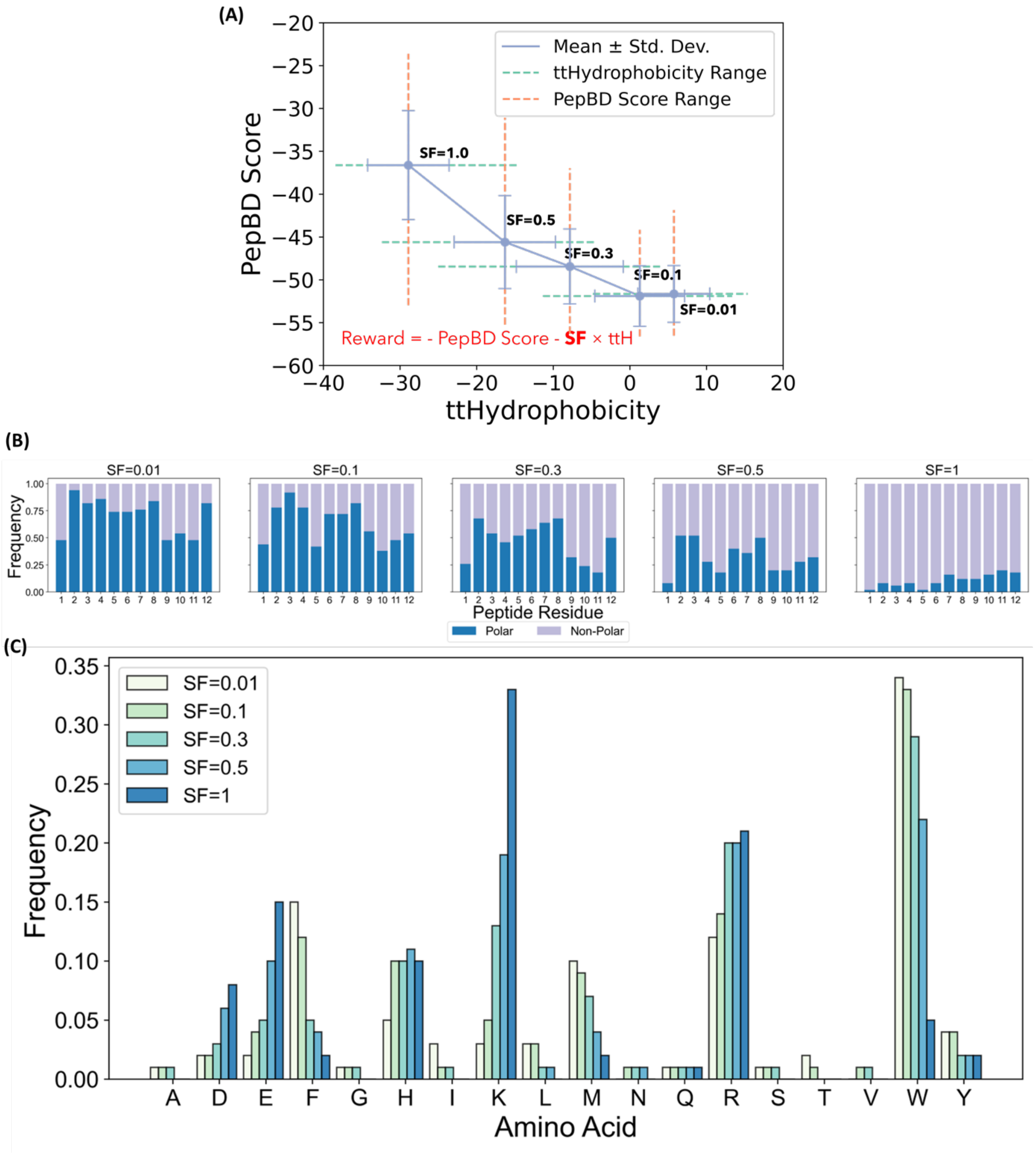


**Figure S5. Simultaneous optimization PepBD score and solubility using the transmembrane tendency hydrophobicity (ttH) solubility score.** A) Pareto optimal solutions for simultaneous optimization of PepBD score and ttH score. Standard deviations and ranges of each value are also shown. (B) Residue polar and nonpolar composition at each position in the peptide for SF values of 0.01, 0.1, 0.3, 0.5, and 1, shown from left to right. (C) Amino acid frequencies at all five SF values. Sequence logos of the PBPs for all SF values are provided in Figure S6. Results in (B) and (C) are from 50 PBPs at each SF value.

Figure S5 shows how the introduction of a transmembrane tendency hydrophobicity (ttH) term into the MCTS reward function influences the PBPs discovered by MCTS. The ttH term is calculated by summing the individual amino acid ttH values. The Pareto front in part A of the figure shows that increasing the relative importance of the ttH score generally leads to a decrease in the PepBD score. The frequency of polar and non-polar amino acids at each position in part B of the figure shows increasing the importance of ttH solubility corresponds to an increase in the frequency of polar residues. In contrast to the CamSol score (Figure 2), ttH does not create amphiphilic peptides. The amino acid concentrations in part C of the figure show that as the importance of the ttH score increases, the frequency of bulky hydrophobic amino acids, like tryptophan (W), methionine (M), and phenylalanine (F), decreases while the frequency of charged residues, like lysine (K) and glutamic acid (E) increases.

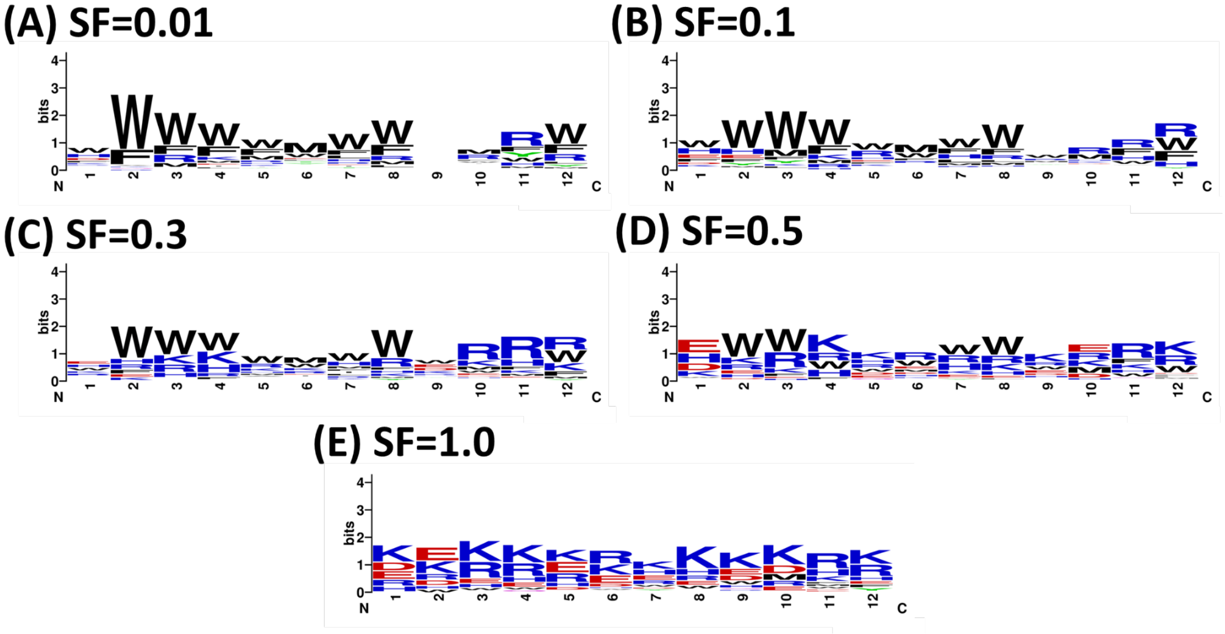


**Figure S6. Sequence logo plots of the PBPs by simultaneously optimizing PepBD score and transmembrane tendency hydrophobicity (ttH) scale at various scaling factor (SF) values.** (A) SF = 0.01. (B) SF = 0.1. (C) SF = 0.3. (D) SF = 0.5. (E) SF = 1.0. SF: scaling factor.

Figure S6 complements Figure S5 by showing sequence logos that visualize where the amino acid types are concentrated in the peptide. To interpret a logo, the size of a letter increases as its frequency of occurring at that position increases. Logos are shown for different SF values to visualize how the amino acid patterns change as the importance of ttH solubility in the MCTS reward function increases (increasing SF increases the importance of the ttH solubility). When SF is small, tryptophan (W) and phenylalanine (F) are common at many positions. As SF increases, the frequency of the hydrophobic residues decreases and the occurrence of arginine (R), glutamic acid (E), and lysine (K) increases. We do not observe a clear sequence motif at any SF value, as indicated by the absence of a common pattern of amino acids in the peptides.

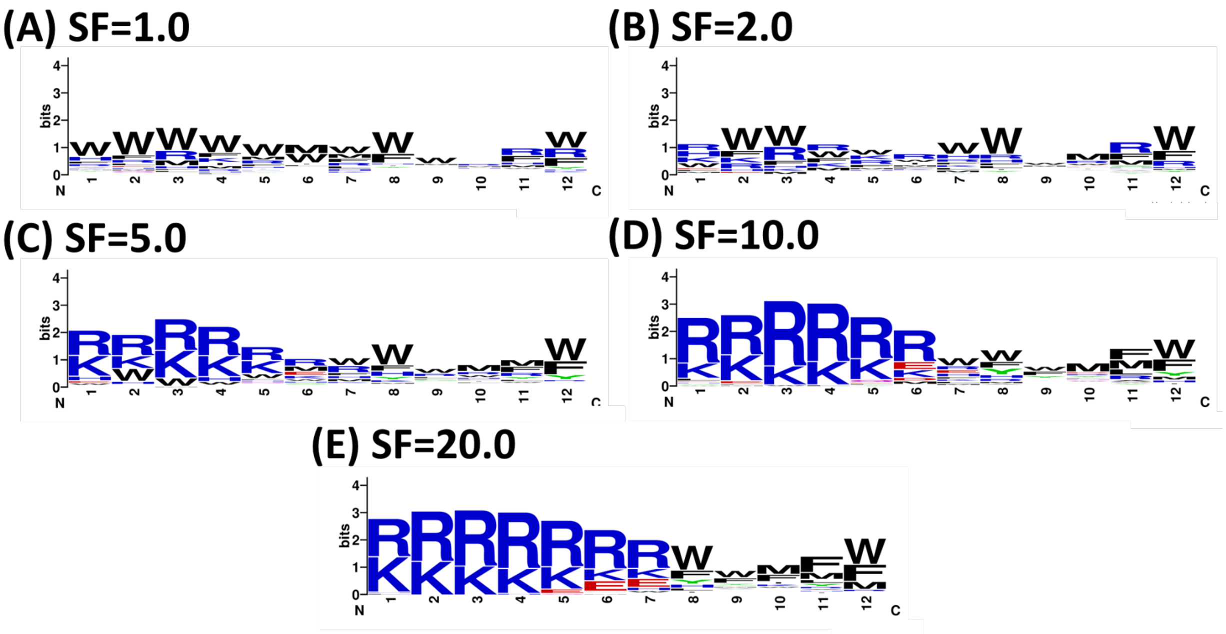


**Figure S7. Sequence logo plots of the PBPs by simultaneously optimizing PepBD score and CamSol with various scaling factor (SF) values.** (A) SF = 1.0. (B) SF = 2.0. (C) SF = 5.0. (D) SF = 10.0. (E) SF = 20.0.

Figure S7 complements Figure 2C by showing sequence logos that visualize how amino acid patterns change as the importance of the CamSol solubility term increases, similar to Figure S6. When SF is small, peptides are enriched in tryptophan (W) and phenylalanine (F) and there is no obvious pattern to the placement of amino acids in the peptide. Increasing SF also increases the frequency of the hydrophobic residues decrease and the occurrence of arginine (R) and lysine (K) increases. As also shown in Figure 2B, hydrophobic and hydrophilic residues are concentrated at the C and N-terminus, respectively, for SF values at 5 or larger.

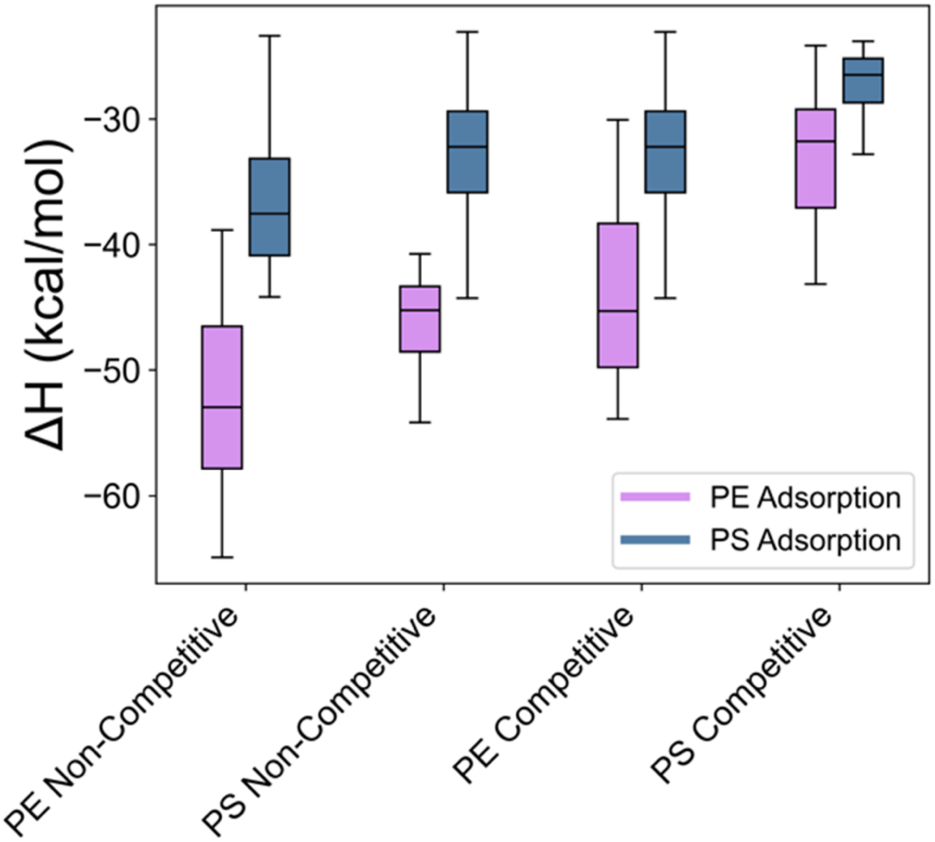


**Figure S8. Binding enthalpies of discovered peptides to polyethylene and polystyrene.** Results are shown for DL PBPs (both competitive and non-competitive discovery) for polyethylene (red) and polystyrene (blue).

Figure S8 shows the binding enthalpies ($\Delta H$) for peptides to polyethylene) or polystyrene (polystyrene). The data complements the binding free energies ($\Delta G$) shown in Figure 4B, where $\Delta G$ is the sum of $\Delta H$ and the binding entropy calculated using normal mode analysis. Results are shown for competitive and non-competitive discovery for both polyethylene- and polystyrene-specific peptides. A peptide is predicted to bind specifically to polyethylene over polystyrene if it has a much more negative $\Delta H$ for polyethylene than polystyrene, and vice versa for a peptide binding specifically to polystyrene. The results in this plot qualitative agree with those shown in Figure 4B, please see discussion associated with the figure for analysis of the plot.

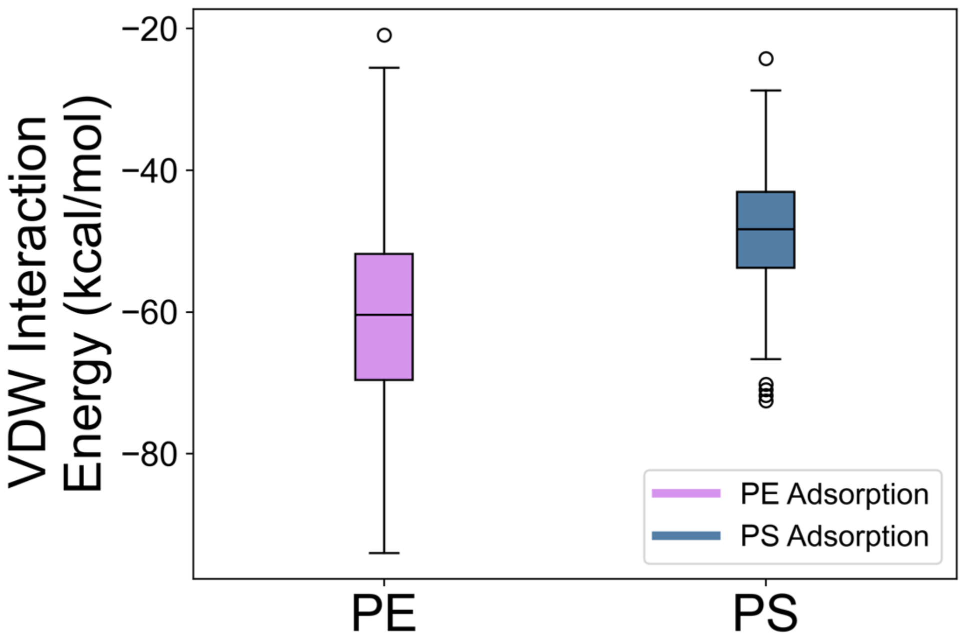


**Figure S9. Distribution of van der Waals (VDW) energies for peptide binding to polyethylene (polyethylene) and polystyrene (polystyrene).** A total of 384 VDW interaction energies are shown for 48 peptides each simulated 8 times. The peptides include 12 polyethylene PBPs, 12 polystyrene PBPs, 12 competitive polyethylene PBPs, and 12 competitive polystyrene PBPs, see Figure 4.

Figure S9 shows the distribution of van der Waals (VDW) interaction energies between the peptide and either a crystalline E or polystyrene surface. Results are for the DL PBPs discovered using CamSol with the scaling factor at 2.0. A total of 384 (48 PBPs × 8 evaluations per PBP) are shown for each plastic. The peptide consistently has more negative (i.e. stronger) van der Waals interactions with polyethylene than polystyrene, which parallels the consistently lower affinity binding free energies of peptides to polystyrene than polyethylene shown in Figure 4B. We hypothesize the van der Waals interaction energies are weaker to polystyrene than polyethylene as the polystyrene surface in MD simulations has more roughness than the polyethylene surface, and the peptide cannot pack as well against a rough surface.

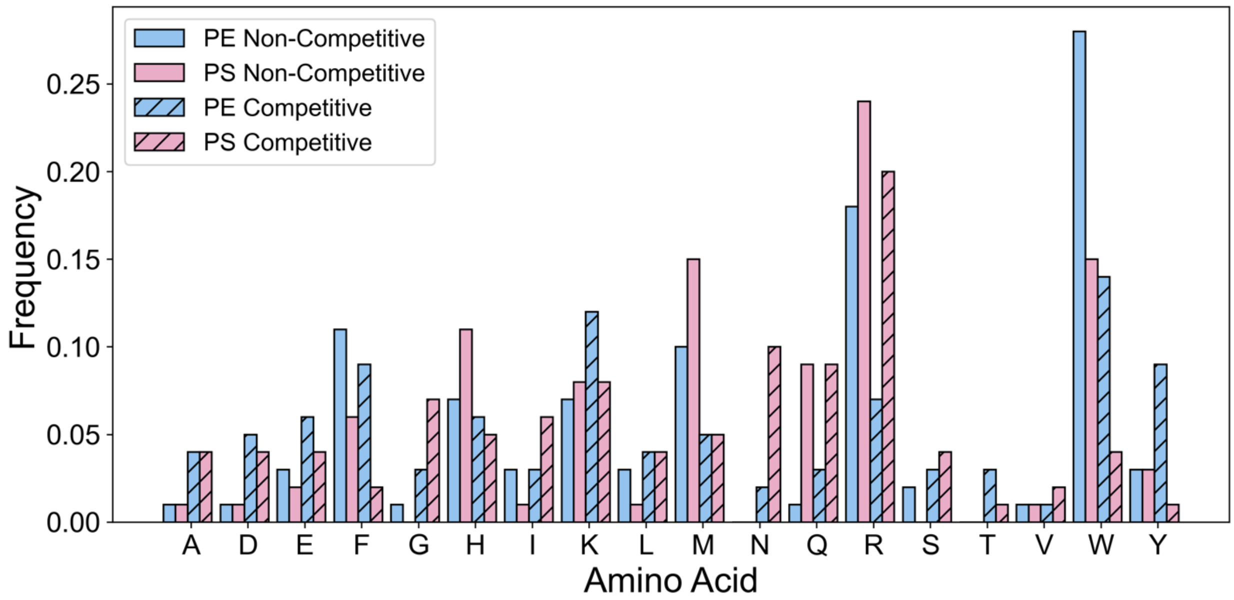


**Figure S10. Comparison of amino acid frequencies using competitive and non-competitive discovery.** Results are shown for 50 DL PBPs for both polyethylene and polystyrene using the competitive and non-competitive discovery methods.

Figure S10 compares amino acid frequencies for PBPs to polyethylene and polystyrene using non-competitive and competitive discovery (see Methods for details on these discovery types). The frequency of some amino acids differs noticeably both between plastics and between competitive and non-competitive discovery. The frequency of tryptophan (W), methionine (M), and phenylalanine (F) is greater in polyethylene PBPs and is lower in competitive discovery. Asparagine (N) is amplified in competitive PBPs for polystyrene, while glutamine (Q) has high frequency for non-competitive and competitive PBPs for polystyrene. Changes in amino acid frequencies indicate that certain amino acids are associated with better binding to one plastic over the other.

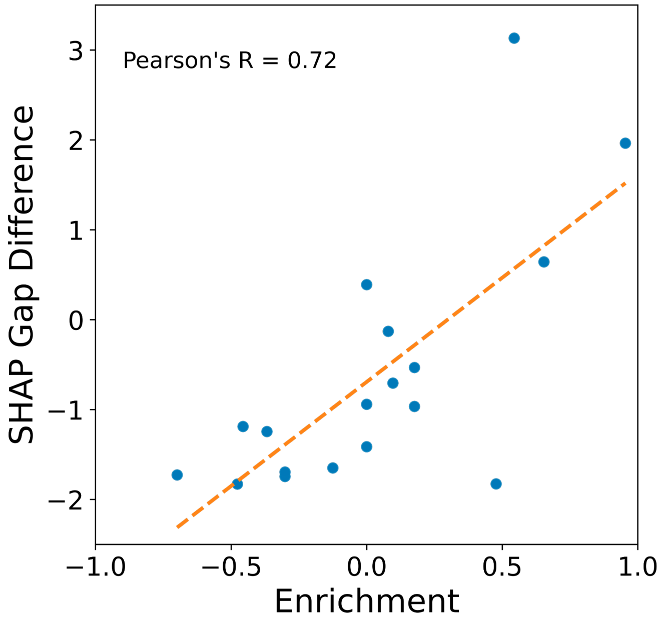


**Figure S11. Correlation between enrichment for amino acid types in competitive discovery (polyethylene versus polystyrene) and the difference in the SHAP Gap values for polyethylene- and polystyrene-specific PBPs.** A SHAP gap value is defined as the difference in the average SHAP value for a set of discovered peptides binding to polyethylene versus polystyrene (Figure 8).

Figure S11 shows a strong positive correlation between the SHAP gap difference and the amino acid enrichment in competitive discovery (polyethylene vs. polystyrene) with a Pearson's coefficient of correlation of 0.72.

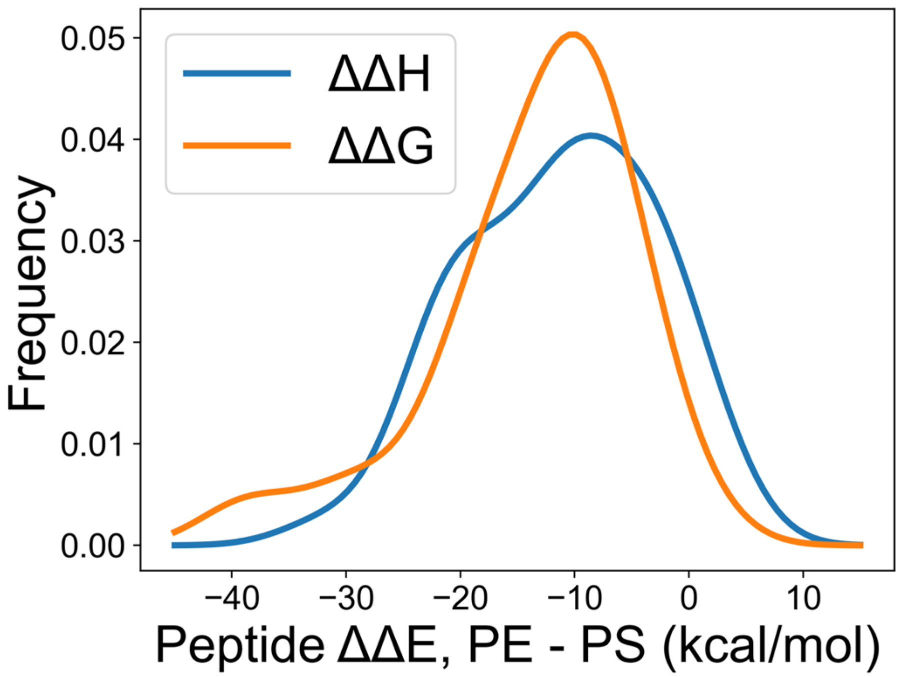


**Figure S12. Distribution of difference in binding enthalpy and free energy between polyethylene and polystyrene peptides.** Data fit with a Gaussian kernel density estimate with a bandwidth of 4. A total of 48 peptides were evaluated to give the fit: 12 polyethylene PBPs, 12 polystyrene PBPs, 12 competitive polyethylene PBPs, and 12 competitive polystyrene PBPs.

Figure S12 combines MD results in Figure 4B and Figure S10 for all discovery types to give the distribution of binding enthalpies and free energies for a total of 48 peptides on both surfaces. Binding enthalpies and free energies were evaluated in MD simulations (see Methods). Peptides on the left tail of the distribution may show much higher relative preference to bind to polyethylene, while peptides on the right tail of the distribution may show greater relative preference to bind to polystyrene.

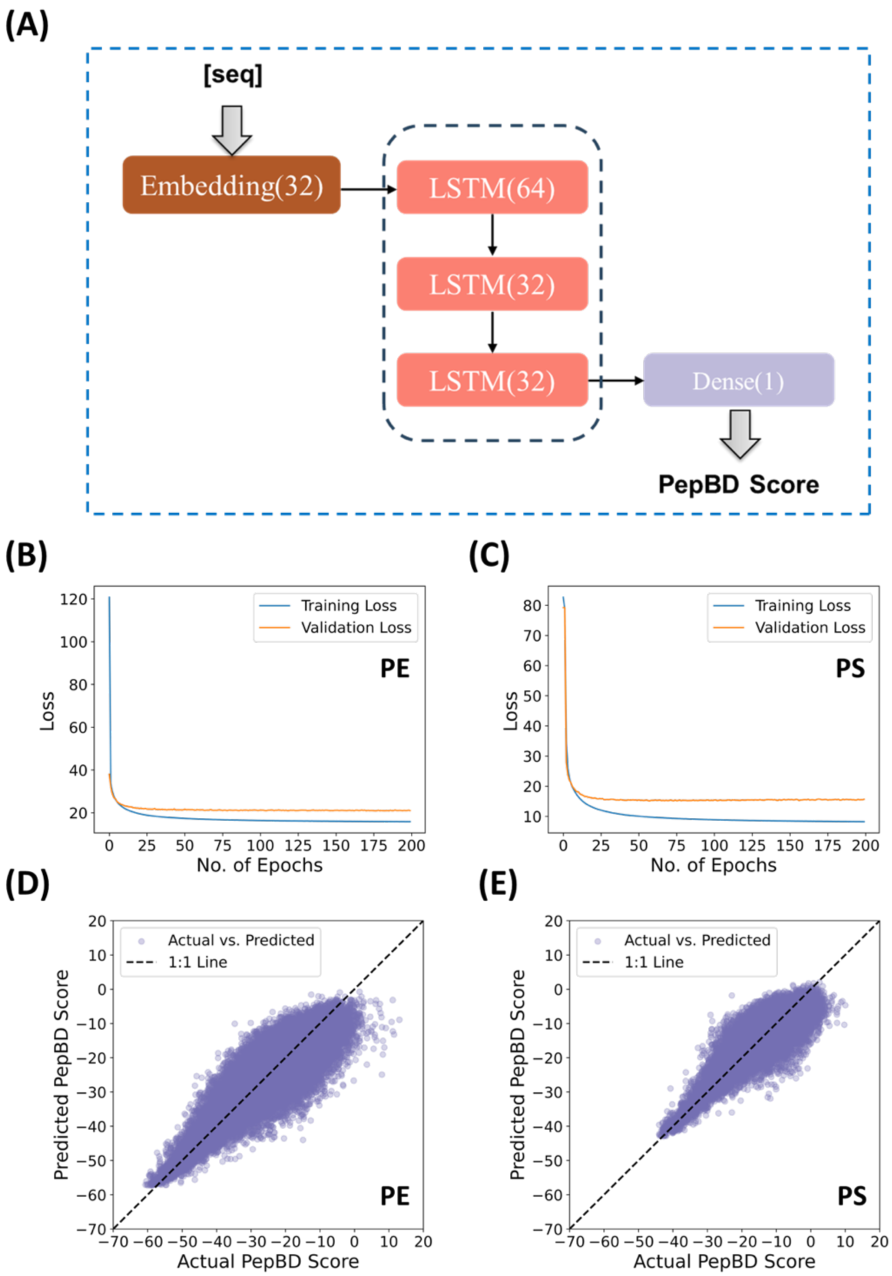


**Figure S13. LSTM architecture, training and validation loss (MSE), and prediction performance on independent test dataset.** (A) The structure of the LSTM model for PepBD score prediction based on a peptide sequence. (B) Learning curves for the polyethylene surrogate model. (C) Learning curves for the polystyrene surrogate model. (D) Predicted PepBD score vs. Actual PepBD score for the 90,107 test data points from the polyethylene dataset. (E) Predicted PepBD score vs. Actual PepBD score for the 40,583 test data points from the polystyrene datasets.

Figure S13 shows the architecture of the LSTM model, which contains one embedding layer, three LSTM layer stacked on top each other, and one dense layer. The embedding layer maps the input sequence to a 32-dimensional vector representation. Subsequent to this stage, the data flows through three stacked LSTM layers, which are configured with 64, 32, and 32 units, respectively. The default hyperbolic tangent function is used as activation in these layers. The dense layer contains one neuron which outputs the predicted PepBD score with the linear activation. The training procedure and prediction efficacy of the surrogate models are also shown for both polyethylene and polystyrene. The polyethylene or polystyrene dataset was partitioned into training, validation, and test sets with respective proportions of 80%, 10%, and 10%. MSE was employed as the criterion for the loss function. Both LSTM models underwent training over 200 epochs, with both training and validation losses plateauing after approximately 100 epochs. This plateau suggests that extending the number of training epochs would unlikely yield significant enhancements in prediction accuracy. The model demonstrating the minimum validation loss during training was chosen for assessing regression performance on the independent test set. The selected models exhibited high accuracy, with the polyethylene model achieving an $R^2$ of 0.82 and an RMSE of 4.58, and the polystyrene model achieving an $R^2$ of 0.81 and an RMSE of 3.90.

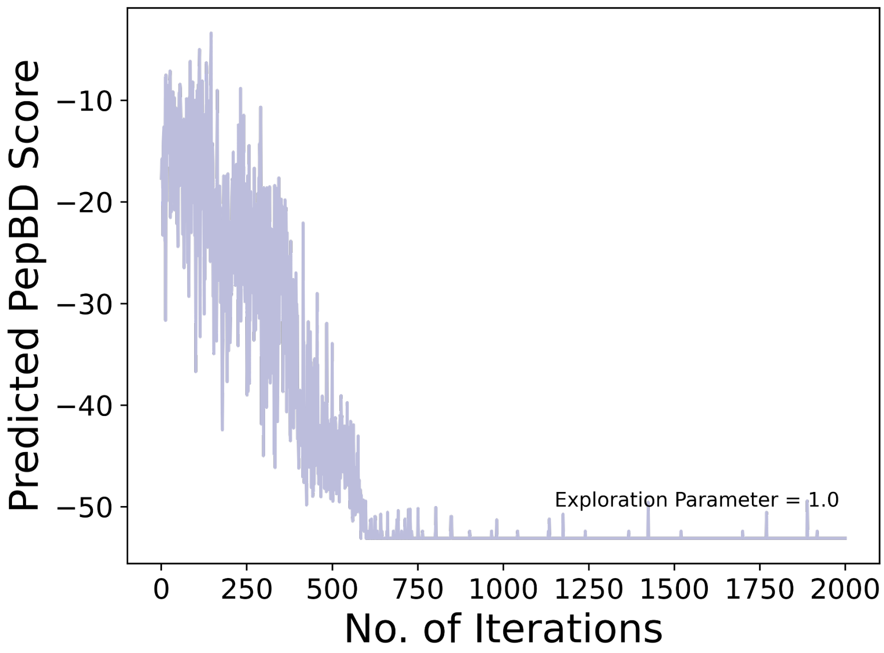


**Figure S14. Predicted PepBD score over the course of 2,000 iterations with an exploration parameter (C) of 1.0 for generating one new peptide.**

Figure S14 shows the optimization process of the DL framework for generating one new peptide. The purple curve tracks the predicted PepBD score of the generated peptide over the loops of MCTS operation. It shows a steep decreasing trend during the first 500 iterations, suggesting that the MCTS optimizes the predicted PepBD score of the sequences generated through prioritizing exploration at this stage. After that, the curve reaches a plateau that is stabilized till the end of the iterations with a predicted PepBD score of approximately –50.
